# A high-throughput method to computationally develop candidate adverse outcome pathways in humans: a proof of concept with insecticides and Parkinson Disease

**DOI:** 10.64898/2026.05.31.728726

**Authors:** Dorian Rollin, Chenyu Shen, Ksenia Groh, Marissa B. Kosnik

## Abstract

Adverse outcome pathways (AOPs) describe stressor non-specific sequences of events between a first molecular trigger (molecular initiating event, MIE), causally linked key events (KEs), and an adverse outcome (AO). AOPs are intended to aid in chemical toxicity testing as a new approach methodology. However, commonly used AOP development methods depend on manual curation, which is labor intensive. As a result, there are still relatively few AOPs and a huge number of toxicity mechanisms and possible adverse outcomes remain undescribed. Therefore, systematic and high-throughput approaches to predict new AOPs are needed. Here, we developed and implemented a data integration-based framework to generate new candidate AOPs using insecticides and Parkinson Disease as a proof of concept. We integrated and statistically linked disconnected databases (e.g., Comparative Toxicogenomics Database, Human Protein Atlas, and Gene Ontology) to form MIE – KE (cell level) – KE (tissue level) – AO candidate AOPs. Through this systematic process, we generated 562,117 candidate AOPs, which we then scored using a weight of evidence (WoE) approach and prioritized 12,756 AOPs with a WoE >0.5. Through random sampling of 100 prioritized AOPs, we found 70% had external literature supporting their biological plausibility, and only 15% represented identifiably implausible associations. The prioritized AOPs describe varied mechanisms of toxicity related to e.g., MAPK, PTEN, and FGFR signaling pathways, with “increases phosphorylation of *MAPK1*” as the most frequent MIE. Our AOP generating approach yields consistently structured AOPs and can complement existing and emerging development methods to expand AOP coverage across different stressors and outcomes.

## 1. Introduction

It is estimated that there are over 350,000 chemicals registered for use globally (Wang et al. 2020), and the majority of these chemicals have not been adequately tested for their potential human health effects. New approach methodologies (NAMs) that harness in vitro and in silico methods have been proposed by regulatory agencies globally to address the challenge of assessing so many chemicals for the plethora of potential adverse outcomes (Krewski et al. 2020). NAMs may also better account for human health effects given the growing evidence that in vivo studies with rodent models do not adequately protect human health (Mak et al. 2014). To aid these efforts, the adverse outcome pathway (AOP) concept was proposed as a knowledge organization tool to describe stressor non-specific pathways linking exposures and adverse effects (Ankley et al. 2009). AOPs comprise the sequential chain of key events (KEs) between the first trigger of an adverse outcome (a specialized key event called the molecular initiating event, MIE) and the adverse outcome itself (AO), with the linkages between points in the chain (e.g., MIE – KE, KE – KE) termed key event relationships (KERs). The AOP framework can be used to organize mechanistic information describing the role of stressors in development of AOs and is of interest to multiple regulatory agencies globally (e.g., the US Environmental Protection Agency (Thomas et al. 2019) and OECD (OECD 2017). For example, by identifying molecular processes in toxicity pathways within an AOP, molecular perturbations measured with high-throughput assays can be linked to apical endpoints used in decision making. Therefore, AOPs are a promising approach to characterize potential health effects for numerous chemicals without reliance on animal testing. However, the process for developing, validating, and incorporating AOPs into AOP databases (e.g., the AOP-Wiki https://aopwiki.org/) is slow and depends on extensive manual efforts. Because of this, there have been calls for the AOP development process to be more pragmatically approached in order to increase the pace of development and regulatory acceptance (Svingen et al. 2021).

There have been several novel approaches for developing individual AOPs using e.g., omics data or causal networks (Azimzadeh et al. 2022; Perkins et al. 2022; Ramšak et al. 2022; Chen et al. 2025). The pace of AOP generation could also be increased through e.g., data integration and text mining (Oki et al. 2016), and tools like AOP-helpFinder have been developed to aid in the process (Jaylet et al. 2025). However, most data integration efforts to date have focused on linking existing AOPs to external information to improve AOP interpretation and application (Mortensen et al. 2021; Saarimäki et al. 2023). Thus, a limitation of current high-throughput tools has been their reliance on the already-existing AOP information. Due to this, the number of existing AOPs is still small (581 in the AOP Wiki as of May 2026), meaning that the spectrum of toxicity mechanisms describing all potential stressor – AO relationships could still be inadequately represented. Additionally, despite calls for standardization, AOPs tend to be developed independently by individual researchers, each using their own structure and vocabulary for describing steps in the AOP. Because of this, there is insufficient consistency in the levels of biological organization represented in AOPs, the way KEs are described, and the corresponding information available for each AOP. Therefore, more high-throughput and standardized methods for AOP development are necessary to harness the full potential of this organizational framework.

We hypothesize that by broadly integrating disconnected datasets that cover different levels of biological organization within the AOP structure, new candidate AOPs can be developed in a systematic and high-throughput manner. In so doing, we propose a workflow that results in a standardized structure, vocabulary, and consistent information being present in each generated AOP. As a proof-of-concept, we focused on insecticides and Parkinson Disease as there is both epidemiological and experimental evidence of a relationship between insecticide exposure and Parkinson Disease development in humans (Höllerhage 2025) and both pesticides and Parkinson Disease contain ample starting data to demonstrate the power of this AOP development approach.

## 2. Methods

A simplified overview of the AOP development process and target structure is shown in Figure 1. Each KE is associated to a level of biological organization according to the example provided in the OECD guidance for AOP development (OECD 2017). A full overview of the steps in the AOP generation process is in SI Figure S1.

**Figure 1.**
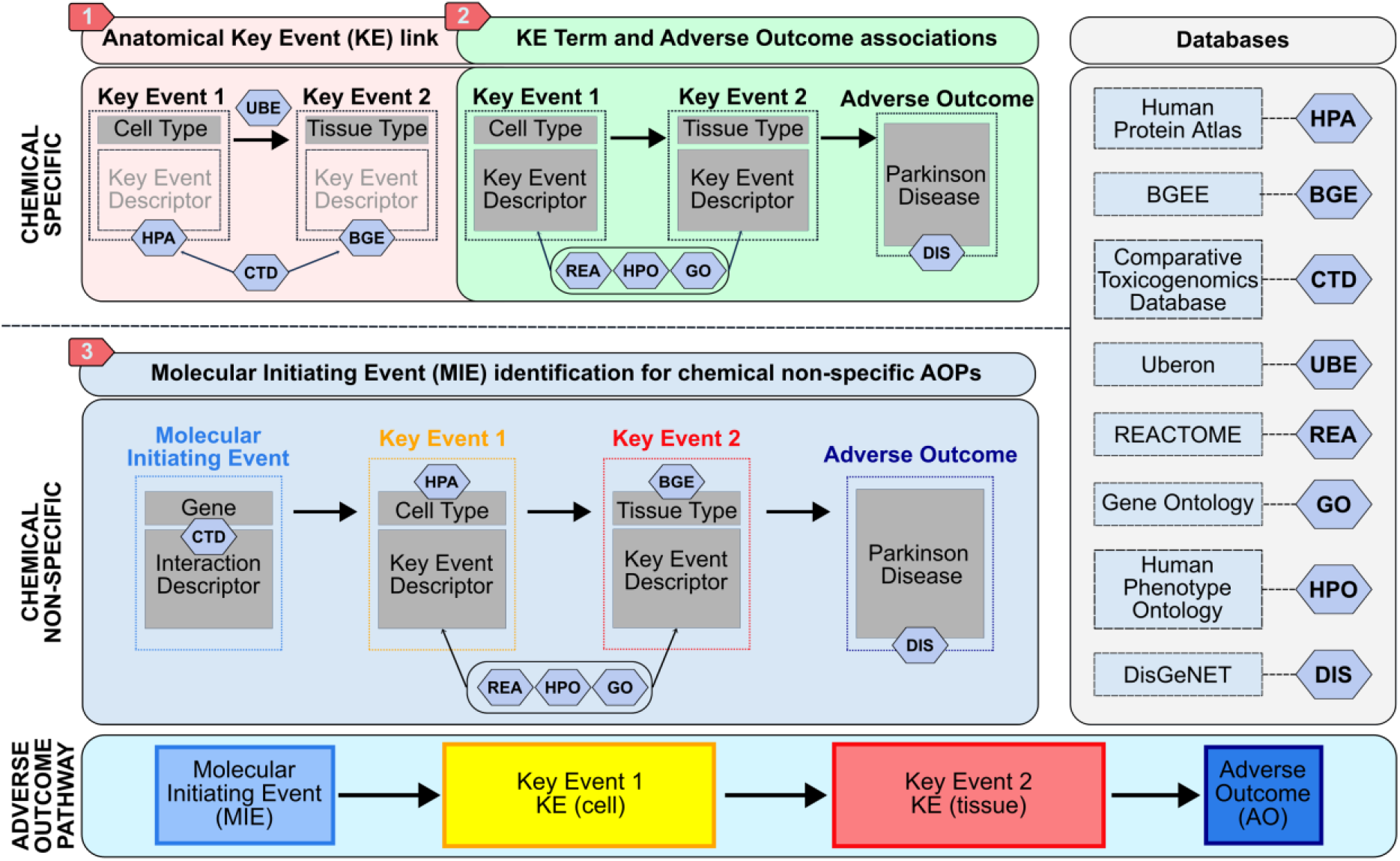
An overview of the AOP development process implemented in this work. Chemical-specific data integration and key event (KE) linkage formation processes (1, 2) followed by identification of a molecular initiating event (MIE) through a chemical non-specific process (3) result in the development of a candidate AOP adhering to a chemical non-specific target structure shown in the bottom panel.

### 2.1. Data curation

The databases used to form the AOPs are shown in SI Table S1. Briefly, curated chemical-gene interactions were downloaded from the Comparative Toxicogenomics Database (CTD) and filtered to contain only *Homo sapiens* (Davis et al. 2025). This database contains curated linkages from the literature. For each chemical in the dataset, one or several interacting genes are linked with one or several interaction types (e.g., Chemical A increases activity of Gene B). Chemicals were reduced to insecticides listed in the Pesticide Properties DataBase (PPDB) using CAS registry numbers for matching and associated with between 5 – 1000 genes (to remove poorly annotated or comparatively strongly annotated chemicals (range from 1 – 4,692 genes per chemical in the starting dataset with a median of 7 genes)). After this filtering, 118 insecticides remained in the chemical-gene interaction dataset from CTD.

Molecular function, biological process, and cellular component Gene Ontology (GO) associations for *Homo sapiens* were collected (Ashburner et al. 2000; The Gene Ontology Consortium et al. 2023) for KE terms. This dataset was filtered using the set of human protein-coding genes available on the *Gene* dataset of National Library of Medicine (NCBI) to remove non-coding, tRNA, rRNA related genes as well as pseudogenes or specific locus names (O’Leary et al. 2024). REACTOME pathways were also collected. This resource contains manually curated and peer-reviewed pathways information with genes known to be associated (Milacic et al. 2024). Human phenotype data (corresponding to the Human Phenotype Ontology, HPO), downloaded from the MONARCH Initiative (Gargano et al. 2024; Putman et al. 2024), was also included for KE term identification.

Tissue type expression was downloaded from Bgee, a database developed and maintained by the *Swiss Institute of Bioinformatics* (Bastian et al. 2025). Bgee gathers data from various studies about gene expression within tissues. By employing a ranking/weighting system normalized across each data type, Bgee generates an expression score used to prioritize the genes that are the most expressed in a specific anatomical entity. As before, filtering using the human gene set of NCBI retained only protein-coding genes.

Cell type expression data was downloaded from the Human Protein Atlas (HPA) (Karlsson et al. 2021). HPA contains the expression in Transcript per Million (TpM) of human protein-coding genes for 85 cell types from 36 different tissues. Using those values, a gene ranking was generated based on the expression score method provided by Bgee to prioritize genes within cell types and apply an equivalent scoring approach at both the cell and tissue type levels. For each anatomical structure (cell or tissue type), only expression scores of 80 or more per gene were retained per biological system to keep only the genes highly expressed within the anatomical structure (starting range from 0 – 100).

Human anatomical ontologies were gathered from Uberon to identify links between the cell types of HPA and the tissue types of Bgee (Mungall et al. 2012). The ontologies include information from Cell Ontology and can, therefore, be associated to the correspondent cell type in HPA (Diehl et al. 2016). Uberon describes tissue-tissue and tissue-cell type associations with “parent-child” relationships, meaning that one tissue type can contain one or several sub-tissues or cell types, and each sub-tissue may also contain sub-tissues or cell types. Bgee uses the Uberon anatomical structures identifiers natively to specify the location of gene expression.

Gene-Disease relationships were downloaded from DisGeNET (Piñero et al. 2020). Gene-Disease data were filtered to the disease “Parkinson Disease”. The distribution of genes per starting data type is in SI Figure S2.

### 2.2. Components of AOP backbone

The AOP backbone was defined as the cell type – tissue type anatomical linkage to Parkinson Disease.

#### 2.2.1. Cell and tissue components

UBERON was used to define the connection between the cell types and tissue types if a tissue (parent) possessed one of the cell types (child in the tree structure). As Bgee is already using Uberon’s IDs for tissue name, only a mapping of HPA IDs was necessary. When available in the tree-structure, HPA IDs were kept for the cell type – tissue type anatomical linkage. Otherwise, an equivalent tissue type was manually identified in the available IDs of Uberon when a similar name was available (e.g., cell types related to “skin” were linked with the Uberon entry “UBERON_0002097” = “skin of body”). Through this, we selected the Bgee tissues that were direct descendants of the “brain” tissue in the tree structure of Uberon, and the corresponding six cell types.

#### 2.2.2. Adverse outcome

Genes known to be implicated in Parkinson Disease were gathered from DisGeNET to generate a gene set for the AO. Overrepresentation analysis was conducted between the tissue type gene set and Parkinson Disease set with the total number of human protein-coding genes as the background (20076 genes), and significance set as p < 0.05 after multiple test corrections with false discovery rate. Six tissues were selected to match the number of cell types, and these were selected from the tissue types most significantly associated with Parkinson Disease and having the most associated genes.

### 2.3. Construction of chemical-specific KE linkages

#### 2.3.1. Development of KE descriptor lists for cells and tissues

To describe what happens at the cell and tissue level in the AOP, candidate KE descriptors were identified from GO, HPO, and REACTOME terms (here forth called KE terms). However, because these data are intended to describe a wide variety of general biological processes, the starting datasets contained many terms that were either too generalized or not relevant for our purposes. To filter the full KE terms list to terms that would be meaningful as KE descriptors, terms were filtered according to three criteria. First, all KE terms that did not describe clear activity were removed. This was done by building a vocabulary describing activity processes, starting from CTD interaction terms included in the Chemical – Gene interaction dataset (e.g., “affects”, “reacts”, “oxidation”). The endings of each term were generalized (e.g., “affects” becomes “affect”, “oxidation” becomes “oxidat”) and the list was appended with three additional directional descriptors that we found commonly implicated across KE terms (i.e., “positive”, “negative”, and “regulat” derived from “regulation”), resulting in a set of 59 activity process terms (SI Table S2). Applying this set to the starting set of KE terms reduced the number from 50,963 to 24,598, removing generalized terms and retaining activity terms that could be meaningful as a KE in an AOP (e.g., selecting for “negative regulation of apoptotic process” and removing the term “apoptotic process”).

The second filtering was done to remove terms that describe human diseases as these would be more suited as an AO rather than as KE terms. This was done using the CTD disease vocabulary including MeSH and OMIM diseases. This list was appended with three additional terms commonly found across KE terms: “Virus”, “Viral”, and “Cancer”, and the term “Death” was removed from the list (so, e.g., cell death processes were not inadvertently filtered out of the KE terms). This resulted in a set of 8,474 disease terms, and 313 KE terms containing these disease terms were removed (e.g., “regulation of viral life cycle”, “Reactive hypoglycemia”, and “SMAD2/3 Phosphorylation Motif Mutants in Cancer”). The third criterion was that only KE terms with greater than 5 and fewer than 1000 genes were kept. This was done to ensure that terms were sufficiently annotated in the starting dataset to represent meaningful KEs, but did not have so many genes that they would describe general processes (e.g., “Signal Transduction”). This yielded a final set of 4,537 KE terms relevant for cell or tissue types.

To build as consistent a structure across AOPs as possible, KE terms were restricted to either cell locations or tissue locations if there was a strong enrichment across this biological level (using overrepresentation analysis between the cell location genes and KE term genes or the tissue location genes and KE term genes as described above for 2.2.2). Then, for all enriched terms in the cell or tissue type, either the cell location or tissue location was selected based on either a 2+ fold greater number of cells/tissues having a term enriched (e.g., “Folding, assembly and peptide loading of class I MHC” was implicated in 4/6 tissue types, but only 1/6 cell types, so it was assigned to the tissue level) or a p-value being one or more orders of magnitude stronger in cells or tissues (e.g., “ubiquitin protein ligase binding” was implicated in all cell and tissue types, but the average tissue p-value was 4e-10 compared to the cell p-value of 1.8e-7, so it was assigned to the tissue level). If a term appeared with similar strength/frequency in both cell and tissue types, then it was left as a possible KE term in either structure. This process resulted in a set of 578 KE terms assigned to cells, 642 KE terms assigned to tissues, and 330 shared KE terms that could appear in either structure.

#### 2.3.2. Chemical-specific KE (cell) – KE (tissue) – AO linkage development

To assign KE terms to the cell and tissue location as part of an AOP (i.e., to construct KE (cell) – KE (tissue) – AO linkages between the KE terms and the AOP backbone), a set of cell and tissue KEs was developed for each of the 118 chemicals. This was done starting from the AOP backbones developed (methods 2.2) and by determining enriched KE terms (methods 2.3.1) for the cell and tissue level that overlap with chemical-specific gene sets. For each insecticide in our analysis, the set of cell and tissue genes was filtered to only those genes that were affected by the chemical in CTD. Then, cell KE terms, tissue KE terms, and shared KE terms that were still enriched for a cell or tissue were determined using overrepresentation analysis as described in 2.2.2. From the resulting set of linkages, KE – KE linkages were removed if they 1) were only implicated by a single chemical, 2) had the same KE term for both the cell and tissue level, or 3) had the same KE term in the cell and tissue level but with different directionality (e.g., the cell KE term was “positive regulation of stress fiber assembly” and the tissue KE term was “negative regulation of stress fiber assembly”). This produced in a final set of KE (cell) – KE (tissue) – AO linkages per chemical.

### 2.4. Extraction of recurring KE (cell) – KE (tissue) – AO linkages

By definition, and as specified by the OECD guidance, AOPs are meant to be chemical non-specific. Therefore, recurring KE (cell) – KE (tissue) – AO linkages were identified across the chemical specific AOPs produced. Any linkage occurring in fewer than three chemicals was removed.

### 2.5. Candidate MIE determination

#### 2.5.1. Candidate gene selection

Because genes identified in this AOP construction process may not represent the “true” starting point for the subsequent chain of events, we term these “candidate MIEs” with the understanding that they may represent additional KEs and an alternative earlier MIE may be identified for a given AOP in the future. For each KE (cell) – KE (tissue) – AO linkage generated in 2.4, genes implicated in the cell level KE across the chemicals generating the candidate AOP were gathered and considered to represent candidate MIEs. This dataset was then merged with the list of genes directly associated with Parkinson Disease (as defined in DisGeNET). However, genes not directly associated with Parkinson Disease were not removed as they may be relevant in Parkinson Disease progression, but not directly annotated in DisGeNET (e.g., *BCL2*).

For the MIE determination, candidate MIE genes per KE (cell) – KE (tissue) – AO linkage were prioritized based on the number of chemicals implicating a given gene in the linkage and (where possible) by accounting for the strength of the gene’s association with Parkinson Disease based on the DisGeNET Disease Specificity Index (DSI). This metric represents the likelihood of a gene being associated with one or several diseases. Values can range from 0.23 to 1, the latter meaning that the gene is specific to only one disease. Therefore, the higher a gene’s DSI is, the more likely it is specific to Parkinson Disease.

To prioritize the candidate MIEs for each KE (cell) – KE (tissue) – AO linkage, we calculated an “MIEscore” for each gene using the following formula:

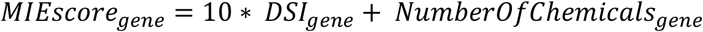

Here, the DSI was taken directly from DisGeNET and multiplied by 10 so that the score is between 0 and 10 rather than 0.23 and 1. The “Number of Chemicals” was the number of insecticides in a recurring KE (cell) – KE (tissue) – AO linkage known to interact with this gene in CTD. If the gene was not associated with Parkinson Disease in DisGeNET and therefore did not have a DSI (e.g., *BCL2*), then just the number of associated chemicals for that KE (cell) – KE (tissue) – AO linkage would determine the MIEscore (as the DSI would be 0). Therefore, this score prioritizes candidate MIEs implicated by many chemicals with additional strength given to genes implicated with specificity in Parkinson Disease. These candidate genes were then ranked according to the MIEscore. Those with an MIEscore > 4 were retained.

#### 2.5.2. MIE descriptor

For each chemical-gene interaction, CTD provides one or several interaction terms (e.g., Chemical A increases activity of Gene B, Chemical A binds to Gene B, etc.). Note that while it is possible the interaction term could refer to a protein rather than a gene, we are keeping the term “gene” here for simplicity as that is the standard term used in CTD. Therefore, for each candidate MIE associated with a KE (cell) – KE (tissue) – AO linkage, the gene would have interaction labels for at least three different chemicals (since a minimum of three chemicals were required for a candidate MIE to be associated with a given KE (cell) – KE (tissue) – AO linkage). For each candidate MIE gene, the top 30% of interaction terms were retained, so long as at least three chemicals had the same interaction term for the candidate gene. If the same MIE was implicated with the same KE (cell) but with different directions (e.g., increases *BCL2* expression and decreases *BCL2* expression), then only the most prevalent MIE direction was selected if there was a 2 or more fold difference. Otherwise, the candidate MIE – KE (cell) connection was removed.

### 2.6. AOP stability determination

To avoid a single chemical driving the AOP predictions and to assess the strength of the AOP association among different insecticides, we conducted an additional step to prioritize the candidate AOPs by subsetting the starting data and regenerating AOPs following a 10x resampling approach. Steps 2.3 – 2.4 were repeated 10 times with each run completed using a random 80% of the starting chemicals. Then, the number of times the same candidate AOP showed up in each subset run was quantified.

### 2.7. Weight of evidence calculations and AOP ranking

To prioritize candidate AOPs, we built a metric incorporating different weights of evidence (WoE).

#### 2.7.1. Literature support for cell KE – tissue KE linkages

Because each KE (cell) – KE (tissue) linkage was formed only based on anatomical location (i.e., a cell type was linked to a tissue type in Uberon), it was possible that some of the KEs linked to each other did not make biological sense. To overcome this limitation, we used AOP-helpFinder 3.0. AOP-helpFinder (AOPhF) is a publicly available tool released in 2019 that enables screening of the scientific literature to extract association information (Carvaillo et al. 2019). Providing a list of candidate KE terms, AOPhF analyzes peer reviewed abstracts in PubMed to predict KERs between the provided terms using text mining and graph theory. The tool is accessible through a web server and provides a scoring system for KER prioritization (Jaylet et al. 2025). The set of cell and tissue KE terms were fed into AOPhF as a search for Event – Event relationships using standard settings in AOPhF (20% reduced search to remove the first part of the abstract, and with 3/4 event match).

The scoring and labelling system of AOPhF can fall in 5 different categories (“Low”, “Quite low”, “Moderate”, “High” and “Very High”) depending on the p-value associated to the KER. All KE – KE support p-values (KE-KE_score_) were then scaled to a score between 0 – 1 depending on the label. These were scored with “Low” AOPhF label receiving a score of 0, “Quite low” scaled from 0.01 – 0.250, “Moderate” scaled from 0.251 – 0.5, “High” scaled from 0.501 – 0.75, and “Very high” scaled from 0.751 – 1 with 1 having the most support for the KER.

#### 2.7.2. Literature support for MIE – KE linkage

The same process was used to search for support between each candidate MIE and KE1 terms (i.e., cell level KE). The AOPhF output for these linkages was represented as p-values according to the same 5 categories in 2.7.1, and all MIE – KE support p-value (MIE-KE_score_) were then scaled the same way with 1 having the most support for the KER.

#### 2.7.3. Significance score calculation

To assess the strength of a given AOP, we took the number of times an AOP appeared in the resampled subsets and three values from the AOP generation process: the number of chemicals implicating the AOP and the overrepresentation analysis p-values for the cell KE term enrichment and the tissue KE term enrichment. These values were then used to quantify the underlying strength of each AOP using the following formula:

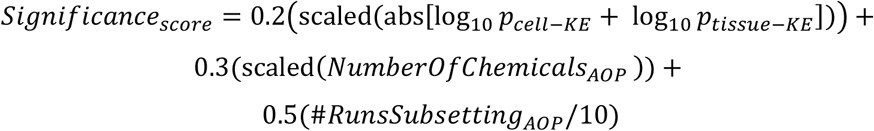

where p is the corrected p-value from each overrepresentation analysis and the absolute sum of the logged p-value scores was rescaled between 0 and 1. The number of chemicals implicating the AOP was also rescaled from 0 to 1 to give comparable weight to this value across AOPs. This score gave the highest weight to a candidate AOP appearing in many of the cross-validation runs (50% of the score) and the potential value for this score was between 0 and 1.

#### 2.7.4. Overall WoE value

The three scores calculated as described in 2.7.1 – 2.7.3 were then combined into a single value as:

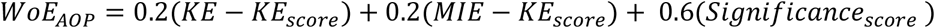

The resulting score between 0 and 1 shows how much support there is for the candidate AOP. Total AOPhF support was given slightly lower weight than the significance score to ensure that associations not explicitly described in the literature (i.e., those with low AOPhF scores) are not significantly down-weighted as these may simply represent associations that have not been well-studied.

### 2.8. Data analysis and visualization

Analyses were performed using R version 4.4.2 (R Core Team 2022). Analyses were conducted using data.table v1.18.2.1 (Barrett et al. 2024), tidyverse v2.0.0 (Wickham et al. 2019), dplyr v1.2.0 (Wickham et al.), tidytext v0.4.3 (Silge and Robinson 2016), scales v1.4.0 (Wickham et al. 2023), stringr v1.6.0 (Wickham 2023), jsonlite v2.0.0 (Ooms 2014), and rentrez v1.2.4 (Winter 2017). Figures were made using ggplot2 v4.0.2 (Wickham 2016), gridExtra v2.3 (Auguie 2017), visNetwork v2.1.4 (Almende B.V. and Contributors and Thieurmel 2025), and treemapify v2.6.0 (Wilkins 2025).

## 3. Results

We generated a set of computationally predicted AOPs by integrating disconnected datasets to form MIE – KE (cell) – KE (tissue) – AO linkages between insecticides and Parkinson Disease. A full walkthrough of this process is shown in SI Figure S1.

First, all possible cell – tissue linkages to Parkinson Disease were determined. This was termed the AOP backbone and was defined using UBERON to assign a candidate Key Event Relationship (KER) between KE1 (cell location) and KE2 (tissue location). This resulted in six cell types linked to 66 tissues. Gene sets of tissues from each AOP backbone were then compared with the gene set of Parkinson Disease using overrepresentation analysis. If the enrichment was significant (p < 0.05), a KER was created between the tissue and the AO Parkinson Disease. To avoid the number of AOPs being inflated by having the same AOP recurring across several different cell – tissue linkages, the top six tissues were selected to match the number of cell types. This yielded AOP backbone links between the cell types “excitatory neurons”, “inhibitory neurons”, “oligodendrocytes”, “oligodendrocyte precursor cells”, “microglial cells”, and “astrocytes”, and the tissue types “prefrontal cortex”, “anterior cingulate cortex”, “CA1 field of hippocampus”, “choroid plexus epithelium”, “inferior olivary complex”, and “right frontal lobe.”

Then, we assigned KE terms to the cell and tissue level for each AOP backbone and insecticide in our analysis. This was done by filtering the cell and tissue level gene sets in the AOP backbone to just those that overlap with a given insecticide. Then, overrepresentation analysis was conducted between the cell level and cell KE terms and the tissue level and tissue KE terms for a given insecticide to identify significant associations (p < 0.05). This resulted in a set of 3,854,821 chemical-specific KE (cell) – KE (tissue) – AO linkages implicated by 71 insecticides. After removing KE (cell) – KE (tissue) linkages implicated by fewer than three chemicals or with conflicting directionality (see Section 2.3.2), the set of KE chains was reduced to 1,480,879. Then, recurring KE (cell) – KE (tissue) – AO linkages were identified across the chemicals, and MIEs were identified based on genes associated with each insecticide for a given KE (cell) – KE (tissue) – AO linkage. Only the top MIEs implicated by at least three insecticides for each AOP were kept. This resulted in a set of 752,225 candidate AOPs, each associated with at least three insecticides. From there, candidate AOPs were removed if they were present as two separate AOPs that were the same but with the directionality different between the two (e.g., the same MIE led to an increase in apoptosis in astrocytes in one AOP and a decrease in apoptosis in astrocytes in another AOP or an increase in activity and a decrease in activity in a given MIE led to the same KE (cell)). While these cases could have biological validity where different conditions or chemical modes of action could cause variable effects, in the interest of being more restrictive than permissive, they were removed. This removed an additional ∼25% of the candidate AOPs.

Our data integration-based approach resulted in 562,117 candidate AOPs with WoE values ranging from 0.004 to 0.77. 12,756 (2%) of the candidate AOPs had a WoE greater than 0.5, and these were selected as the prioritized set of AOPs. Five example AOPs from the prioritized set are shown in Figure 2A. These AOPs demonstrate output with different combinations of the underlying WoE values: the overall top ranked AOP (high AOPhF scores, high significance score), AOPs with either high MIE – KE_scores_ or high KE – KE_scores_ from AOPhF, the AOP with the highest significance score but 0 AOPhF scores, and the lowest ranked AOP that was still included in the priority list. The distribution of significance scores, AOPhF scores, and the resulting WoE values across all AOPs are shown in Figure 2B – E.

**Figure 2.**
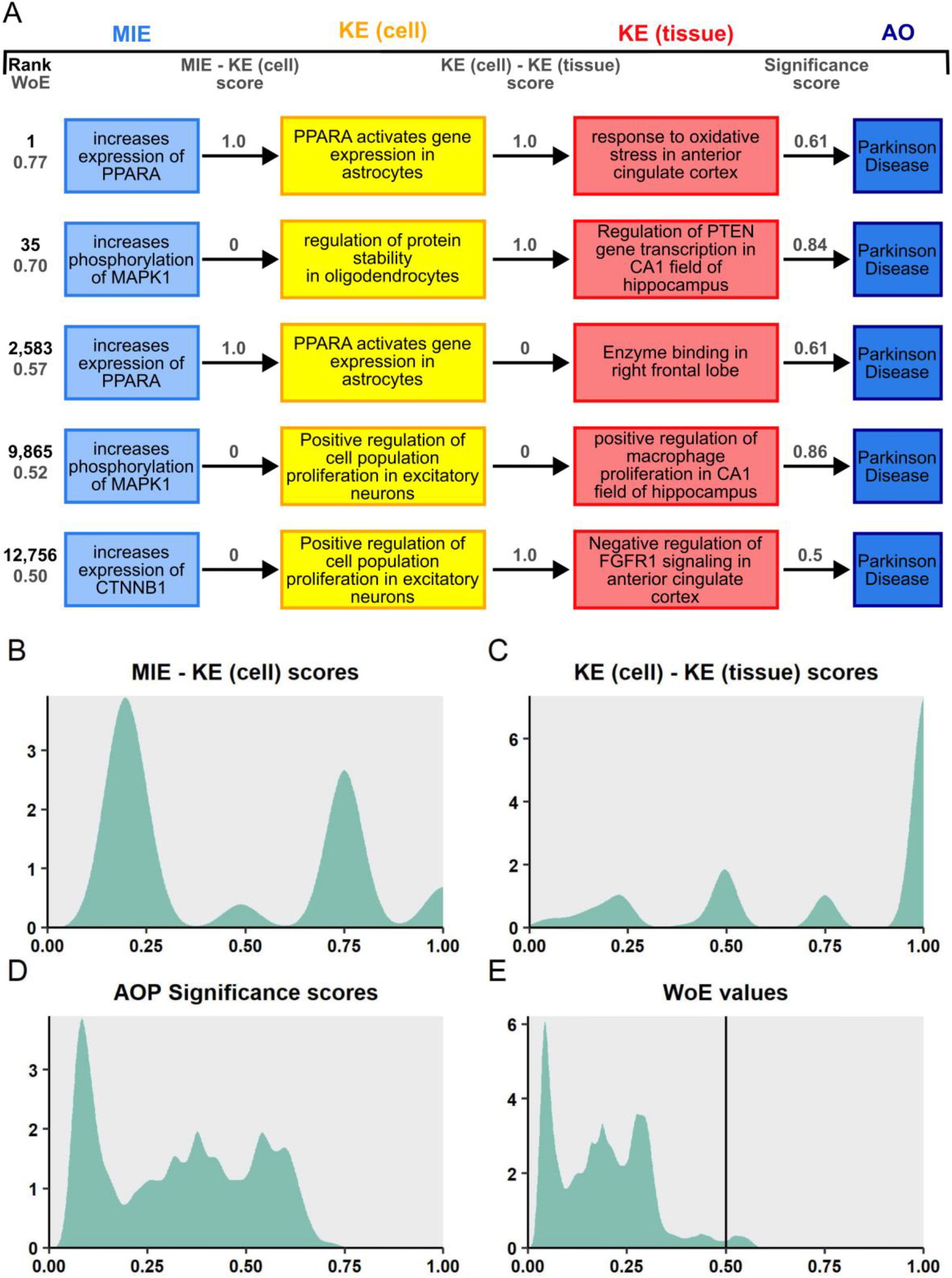
Examples of prioritized AOPs and score distributions underlying the complete set of candidate AOPs. A) Example prioritized AOPs with different underlying AOPhF scores and significance scores: the overall top-ranked AOP, the top-ranked AOP with an MIE – KE (cell) score of 0, the top-ranked AOP with a KE (cell) – KE (tissue) score of 0, the top ranked AOP based on significance score, and the lowest ranked AOP that was still in the prioritized list. Note: while the significance score is shown between the KE (tissue) – AO arrow, this value applies to the whole AOP and is just shown in that location for visualization purposes. B – E) Distributions of the underlying AOP score values across all candidate AOPs (including de-prioritized ones). Only AOPhF scores above 0 are shown. The line in the WoE distribution (panel E) shows the cut-off for the prioritized AOPs.

### 3.1. Top-weighted AOPs implicate similar MIEs and KEs

An overview of the top 500 AOPs is shown in Figure 3. Across these AOPs (WoE 0.61 – 0.77), the KE (cell) “*PPARA* activates gene expression” is the highest-weighted KE (cell), owing to its relevance as both an MIE – KE (cell) link in AOPhF and a KE (cell) – KE (tissue) link in AOPhF (implicated in 89 of the top 500 AOPs). That is, the only candidate AOPs with both MIE – KE_score_ and KE – KE_score_ > 0.8 had “*PPARA* activates gene expression” as the cell KE term (implicated in oligodendrocytes, astrocytes, or microglial cells). Other major KE (cell) terms in the top 500 AOPs include “regulation of protein stability” (61 AOPs), “protein phosphorylation” (74 AOPs), or terms related to protein serine kinase activity (e.g., “protein serine kinase activity” = 120 AOPs, “protein serine/threonine kinase activity” = 120 AOPs). These terms were implicated across all cell types with astrocytes (133 AOPs), microglial cells (97 AOPs), and oligodendrocytes (76 AOPs) as the most common cell types. “Increases expression of *PPARA*” (42 AOPs), “increases activity of *AHR*” (11 AOPs), “decreases expression of *ABCA1*” (38 AOPs), “decreases expression of *BCL2*” (42 AOPs), and “increases phosphorylation of *MAPK1*” (360 AOPs) were the MIEs most frequently implicated. KE (tissue) terms were more variable than the KE (cell) terms, with “regulation of *PTEN* gene transcription” (137 AOPs), “response to oxidative stress” (22 AOPs), terms related to the MAPK pathway (“regulation of stress-activated MAPK cascade”, 53 AOPs; “negative regulation of MAPK pathway”, 56 AOPs), and FGFR signaling terms (“negative regulation of FGFR3 signaling”, 91 AOPs; “negative regulation of FGFR1 signaling”, 76 AOPs) highly implicated. KE (tissue) terms were implicated across all six tissue types with the prefrontal cortex (125 AOPs), anterior cingulate cortex (124 AOPs), CA1 field of hippocampus (121 AOPs), and right frontal lobe (117 AOPs) occurring most frequently.

**Figure 3.**
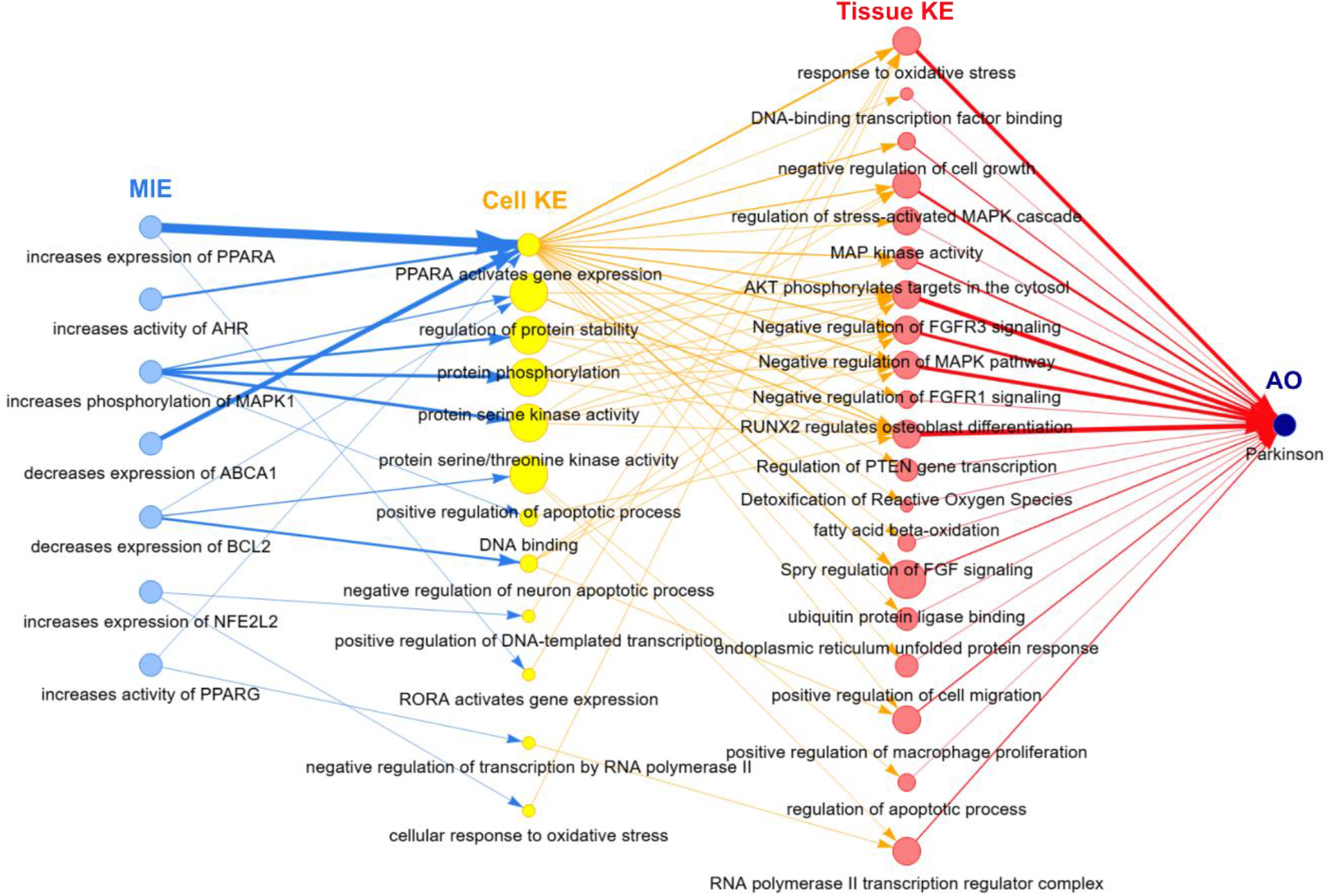
Network of top 500 AOPs constructed based on WoE value, independent of cell or tissue type. The thickness of an arrow reflects how many of the top AOPs include that KER. The size of the KE circles reflects how many cell or tissue types are associated with that KE.

### 3.2. Many recurring KE – KE chains are shared across tissue and cell types

Across all 12,756 prioritized candidate AOPs for insecticide-induced Parkinson Disease (WoE >0.5), there were 54 MIEs, 249 cell KEs (combined cell type and KE term), and 365 tissue KEs (combined tissue type and KE term). 71 total insecticides were associated with at least one AOP with a range of 3 – 17 insecticides associated with each AOP (median of 5).

We found that, across the highest ranked AOPs based on WoE value, a recurring set of KE terms were connected (regardless of cell or tissue), highlighting potentially important generalized mechanisms for further analysis. Figure 4A shows the top 25 KE – KE pairs with the most associated cells and tissues. The most prevalent KE terms overall are shown for cells and tissues in Figure 4B and 4C, respectively.

**Figure 4.**
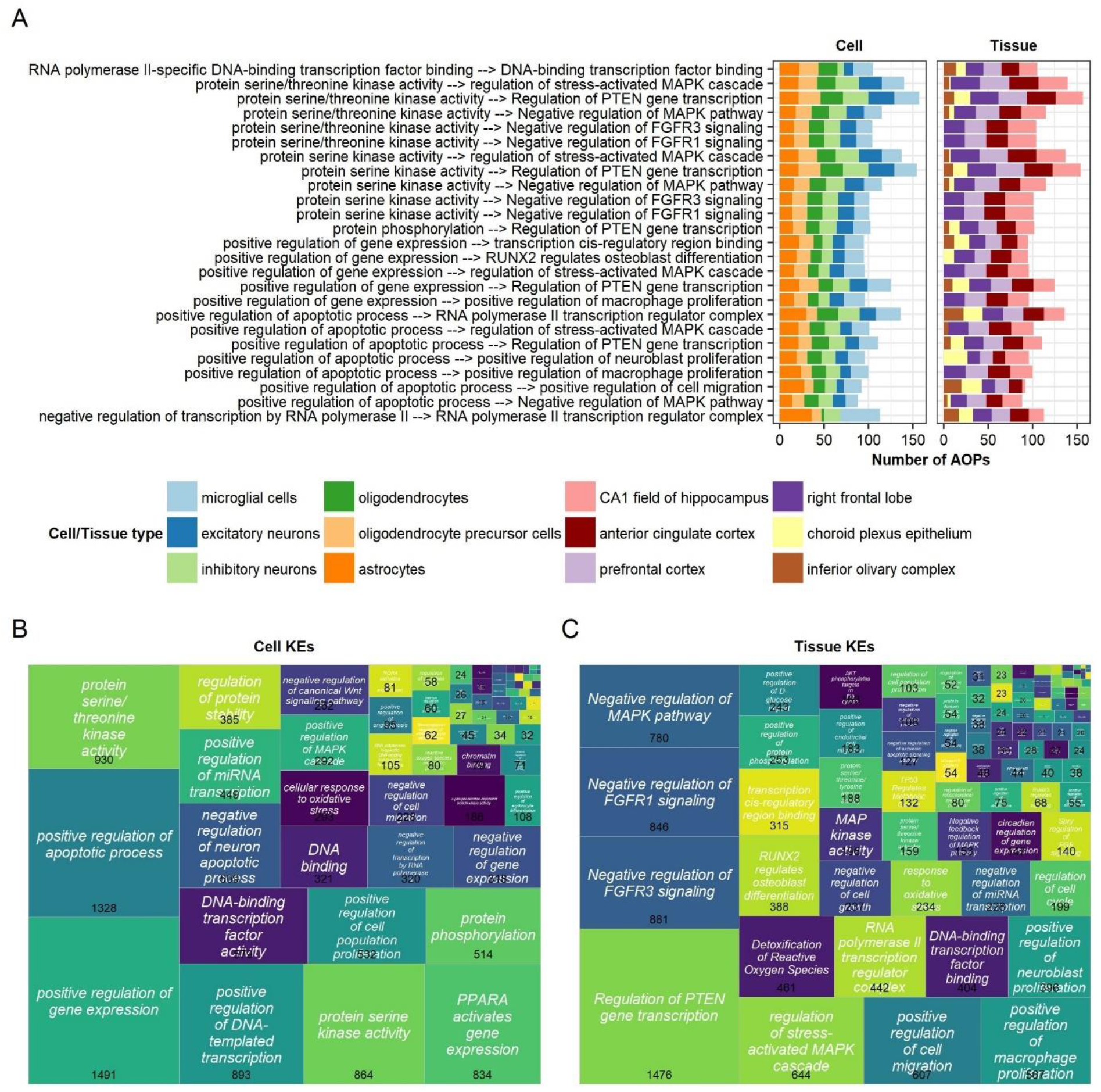
Overview of the KE terms in the prioritized AOPs. A) The 25 most prevalent KE – KE chains recurring across the most different cell and tissue types. B – C) The most prevalent KE term across all AOPs for B) Cells and C) Tissues.

The most frequently recurring KE – KE pairs across all cell and tissue types relate to protein serine/threonine kinase activity or regulation of gene expression as the cell KEs and regulation of the MAPK cascade, *PTEN* gene transcription, and FGFR1/3 signaling as the tissue KEs. Across all prioritized AOPs, the most frequent cell KE terms were “positive regulation of gene expression” (1491 AOPs), “positive regulation of apoptotic process” (1328 AOPs), “protein serine/threonine kinase activity” (930 AOPs), and “positive regulation of DNA-templated transcription” (893 AOPs). The most frequent tissue KE terms were “regulation of *PTEN* gene transcription” (1476 AOPs), “negative regulation of FGFR3 signaling” (881 AOPs), “negative regulation of FGFR1 signaling” (846 AOPs), and “negative regulation of MAPK pathway” (780 AOPs).

### 3.3. Prioritized MIEs include frequent toxicology-relevant genes and Parkinson Disease genes

Disregarding cell or tissue location, there were 1,470 unique MIE – KE term – KE term linkages between insecticides and Parkinson Disease (subset of MIE – KE (cell) linkages in Figure 5). “Increases expression of *NFE2L2*” was the MIE with the most unique MIE – KE term – KE term linkages, particularly in association with “positive regulation of DNA-templated transcription” and “positive regulation of gene expression” contributing to “response to oxidative stress.” “Increases phosphorylation of *MAPK1*” leading to “regulation of protein stability”, “protein serine kinase activity”, and “protein phosphorylation” contributing to “regulation of *PTEN* gene transcription” was another frequent set of linkages. “Increases phosphorylation of *MAPK8*” leading to “positive regulation of gene expression” and “positive regulation of apoptotic process”, and “increases expression of *NFKB1*” leading to “positive regulation of DNA-templated transcription” and “negative regulation of gene expression” are other example chains of MIE – KE (cell) terms that recurred across cell and tissue types. Across all prioritized AOPs, “increases phosphorylation of *MAPK1*” (1,361 AOPs), “increases expression of *NFE2L2*” (1,228 AOPs), “increases phosphorylation of *MAPK8*” (1,205 AOPs), and “increases phosphorylation of *MAPK14*” (1,026 AOPs) were the most frequent MIEs.

**Figure 5.**
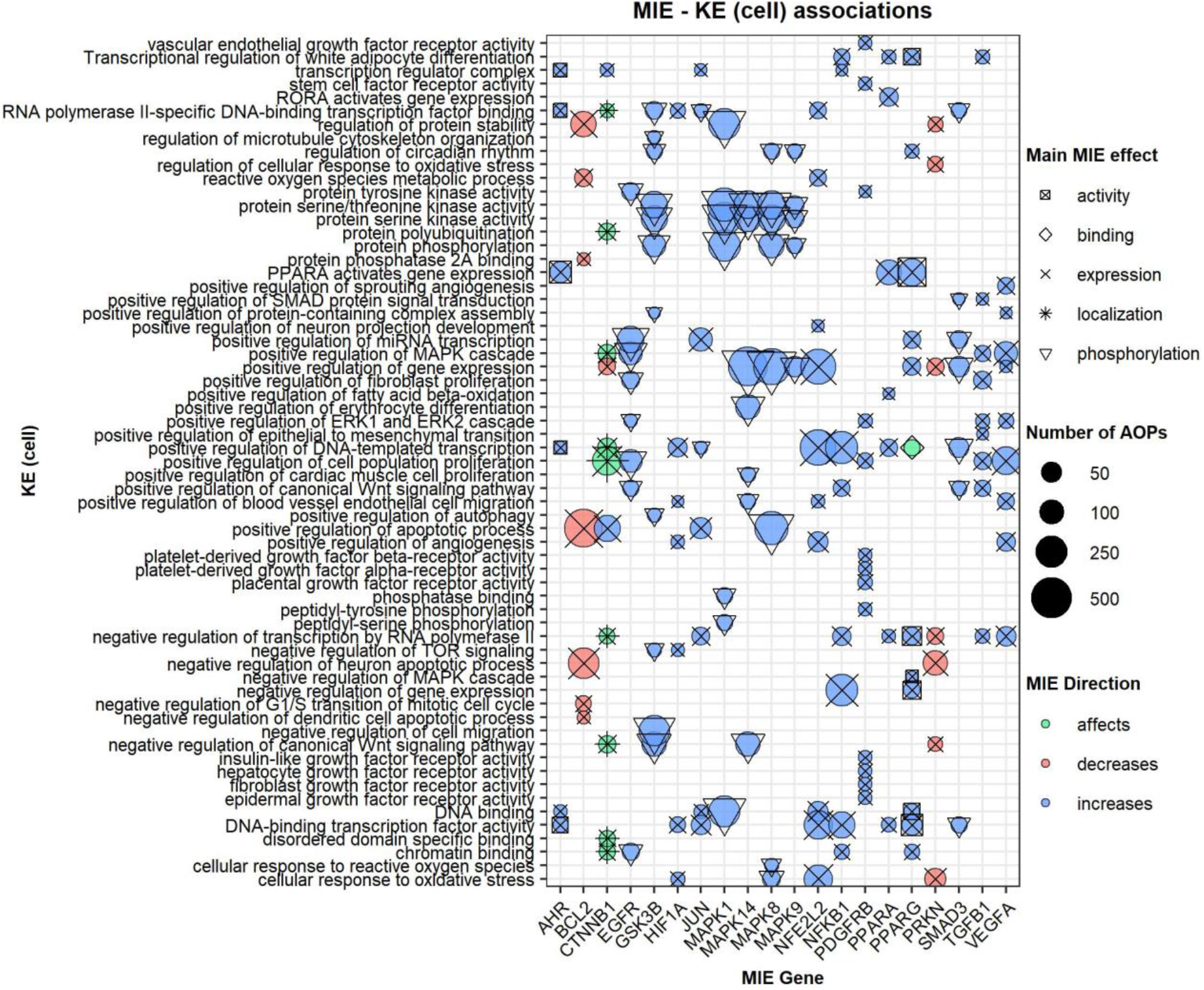
Overview of MIE – KE (cell) associations for the top 20 most frequent MIE genes. The highest weighted MIE activity is shown as the shape overlaying the colored circle which indicates the directionality of the MIE. The size of the bubble shows the number of AOPs (across cell types) with that MIE – KE (cell) association.

## 4. Discussion

We developed a systematic and high-throughput method to generate new candidate AOPs using a data integration approach. With a focus on insecticides and Parkinson Disease, we formed 562,117 total candidate AOPs and prioritized 12,756 AOPs with WoE values >0.5 calculated by combining evidence based on the strength of the full association and the literature support for KERs. Prominent KE terms in these AOPs align with current knowledge of Parkinson Disease. For example, negative regulation of FGFRs was a KE in multiple tissues, and *FGFR1* and *FGFR3* have a crucial role in the central nervous system where they regulate dopaminergic signaling and survival (Grothe and Timmer 2007; Timmer et al. 2007; Wang et al. 2026). An increase in phosphorylation of *MAPK1* was the most frequent MIE, and chronic activation of the MAPK signaling pathway has been observed following prolonged insecticide exposures (e.g., organophosphate and organochlorine insecticides (Farkhondeh et al. 2020; López-Merino et al. 2023)). A decrease in *BCL2* was another prevalent MIE, and, despite not being associated with Parkinson Disease in DisGeNET, this association is implicated in neurodegeneration in Parkinson Disease-induced mouse models (Vila et al. 2001) and is a prominent pesticide target implicated in Parkinson Disease (e.g., for paraquat (Amaral et al. 2025)). Finally, analyzing complete candidate AOPs also reveals biologically plausible associations. The top-ranked AOP describes an increase in *PPARA* gene expression (MIE) leading to activation of *PPARA* dependent gene expression in oligodendrocytes (KE cell) and subsequent occurrence of oxidative stress in the anterior cingulate cortex (KE tissue) preceding Parkinson Disease (AO). This is supported by the literature where studies demonstrated that *PPARA* activation in astrocytes can induce downstream target gene expression (Luo et al. 2023), and that *PPARA* can have an adverse role in astrocytes where agonist-binding can cause *PPARA* activation to induce decreased glutamate uptake through endocytosis of the glutamate transporter *GLT1* in astrocytes (Huang et al. 2017). Dysfunction of such excitatory amino acid transporters is implicated in Parkinson Disease due to an increase in glutamate leading to excitotoxicity, which can trigger oxidative stress and neuronal damage (Zhang et al. 2016; Wu et al. 2025).

While AOPhF can provide MIE – KE and KE – KE support, this depends on whether this relationship has already been reported in the literature. To mitigate this bias, we also incorporated into the final WoE calculation a significance score that describes the statistical strength (based on the enrichment p-values and number of chemicals associated with the AOP) and stability (based on resampling subsets of chemicals and regenerating AOPs) of the candidate AOPs we formed. We find that prioritized AOPs without AOPhF support for both KERs still resulted in biologically plausible associations. For example, the AOP with the highest significance score (0.86) had AOPhF scores of 0. It describes an increase in phosphorylation of *MAPK1* (MIE) leading to regulation of protein stability in oligodendrocytes (KE cell) which progresses to positive regulation of macrophage proliferation in the CA1 field of the hippocampus (KE tissue). There is evidence of the role of *MAPK1* as a regulator of myelination through phosphorylation of myelin basic protein in oligodendrocytes, and an overly increased activity can adversely affect myelin and axonal integrity, and thus function of the central nervous system (Ishii et al. 2012; Ishii et al. 2016). Myelin debris has been of recent growing interest in Parkinson Disease because it can trigger inflammatory responses (Yang et al. 2023), which lead to macrophage mediated myelin debris clearance; this dynamic is implicated in other nervous system diseases like multiple sclerosis (Kopper and Gensel 2018) and Alzheimer’s Disease (Feiten et al. 2026). Overall, we find good support for the candidate AOPs identified through our approach. We conducted a preliminary literature search for 100 randomly selected prioritized AOPs where we searched for references supporting the KERs, cell and tissue locations, and relevance to insecticide exposures and Parkinson Disease, and found that 70% had plausible biological support (SI Table S3). We found that 9% had complete support for all components in the AOP (“very high” support) with an additional 36% and 25% having “high” or “moderately high” support for the biological plausibility of that AOP, with the primary uncertainty being unclear tissue location or potential feedback loops between the MIE and KEs. 15% of the randomly selected AOPs had “moderate” support but inadequate information for all links in the association, and only 15% were not plausible, primarily due to some KE terms that were not plausible for a given tissue location (e.g., terms related to ossification implicated in brain tissue; “low” to “very low” support).

Compared to other AOP-generating approaches, to our knowledge, ours is the first to integrate disconnected datasets and systematically identify recurring chains of connected KEs. These candidate AOPs can serve as a starting point to define putative AOPs for inclusion into the AOP-Wiki and to identify further research needs. As of May 2026, the AOP-Wiki has one OECD endorsed AOP (AOP 3) and five AOPs with Parkinsonian motor deficits as an AO (AOPs 464, 587, 588, 589, and 593). The MIEs for these AOPs are different from those in our dataset and relate to e.g., activation of metabotropic glutamate receptors and binding of inhibitors to NADH-ubiquinone oxidoreductase (complex I), while KEs describe mitochondrial dysfunction and degeneration of dopaminergic neurons of the nigrostriatal pathway. The lack of direct overlap with our candidate AOPs is likely due to differences in the types of MIEs, KE terms and the AO explicitly targeted in our workflow. Therefore, it can be assumed that our candidate AOPs cover an additional biological space affected by stressors linked to Parkinson Disease. An advantage of our AOPs is that each KE has been annotated with a set of corresponding genes, which can inform pathway-based assessments for toxicity testing. Further, all linkages in our AOPs are built following a standard procedure and directed at biological levels, meaning that KEs present across multiple AOPs can easily be assessed using an AOP network structure. This is a benefit compared to existing AOPs that are individually developed with potentially different terminologies (Yarar et al. 2024).

While a promising starting point for high-throughput AOP development, our approach also has several limitations. For example, because data used for MIEs comes from unrefined chemical – gene linkages found in CTD, some of these associations may not represent the true first trigger of a biological outcome, but rather gene activity that happens downstream of a different MIE trigger. Indeed, while some of the MIEs in candidate AOPs describe e.g., phosphorylation of a target, others only describe changes in gene expression. It is also possible that in some cases, there can be an association between an MIE and a KE, but it would be more typical for the KE to precede an MIE, and the MIE activity tends to trigger the KE only through a feedback loop. Future research can attempt refining the MIE selection process through the addition of e.g., molecular docking data, where available.

Another limitation relates to discerning biological meaningfulness in KE – KE linkages. Because the AOP backbone links the KE (cell) to KE (tissue) using Uberon, the basis of that linkage is the presence of the cell in a tissue rather than a direct link between the cell KE descriptor and the tissue KE descriptor. Without AOPhF providing literature support for that association, we cannot know for sure that those two KE processes are linked. This can for example create a conflict in directionality between the KERs (e.g., a decrease of the same gene leads to an increase in apoptosis in one candidate AOP but a decrease in apoptosis in another candidate AOP). While these findings could represent true mechanisms where directionality varies depending on different conditions, we chose to exclude these instances from the current dataset due to the inability to distinguish biologically meaningful cases. Finally, the location of KEs were not considered in AOPhF support, meaning that, even when KE – KE term linkages have support, the biological location relevance may not be fully supported (e.g., as seen with osteoblast differentiation implicated in brain tissue in SI Table S3). Future efforts can enhance our approach by comparing our candidate AOPs to omics data measured following chemical exposures (including in different cell lines) to see if the directionality of gene expression for that chemical aligns with the direction of KEs predicted in our putative AOPs. Further, by pairing our approach with additional AOP development methods (e.g., causal discovery, AI-based approaches), the directionality of associations and true MIEs may be better clarified, and the priority list of candidate AOPs can be further refined. Therefore, we expect our data integration-based approach for AOP generation can complement other emerging AOP development methods by systematically expanding the biological space covered by candidate AOPs across diverse stressor sets and adverse outcomes. Because our workflow is well-defined, it can be readily used to generate AOPs covering additional stressors and adverse outcomes of interest, thus supporting the urgent need for a broader use NAMs in chemical toxicity testing.

## 5. Conflict of Interest statement

The authors declare no conflicts of interest.

## 6. Funding

This work has received funding from Novartis Forschungsstiftung as part of the FreeNovation 2024 scheme (FN24-0000000639). Additional funding came from internal Eawag discretionary funding.

## Notes

### Competing Interest Statement

The authors have declared no competing interest.

## Bibliography

Almende B.V. and Contributors, Thieurmel B. 2025. visNetwork: Network Visualization using ‘vis.js’ Library. https://CRAN.R-project.org/package=visNetwork

Amaral L et al. 2025. The neurotoxicity of pesticides: Implications for Parkinson’s disease. Chemosphere. 377:144348. 10.1016/j.chemosphere.2025.144348

Ankley GT et al. 2009. Adverse outcome pathways: A conceptual framework to support ecotoxicology research and risk assessment. Environmental Toxicology and Chemistry. 29(3):730–741. 10.1002/etc.34

Ashburner M et al. 2000. Gene Ontology: tool for the unification of biology. Nat Genet. 25(1):25–29. 10.1038/75556

Auguie B. 2017. gridExtra: Miscellaneous Functions for ‘Grid’ Graphics. https://CRAN.R-project.org/package=gridExtra

Azimzadeh O et al. 2022. Application of radiation omics in the development of adverse outcome pathway networks: an example of radiation-induced cardiovascular disease. International Journal of Radiation Biology. 98(12):1722–1751. 10.1080/09553002.2022.2110325

Barrett T et al. 2024. data.table: Extension of ‘data.frame‘. https://CRAN.R-project.org/package=data.table

Bastian FB et al. 2025. Bgee in 2024: focus on curated single-cell RNA-seq datasets, and query tools. Nucleic Acids Research. 53(D1):D878–D885. 10.1093/nar/gkae1118

Carvaillo J-C, Barouki R, Coumoul X, Audouze K. 2019. Linking Bisphenol S to Adverse Outcome Pathways Using a Combined Text Mining and Systems Biology Approach. Environ Health Perspect. 127(4):047005. 10.1289/EHP4200

Chen Y et al. 2025. Adverse outcome pathway (AOP) framework for predicting toxic mechanisms in E-cigarette-induced lung injury. Ecotoxicology and Environmental Safety. 302:118703. 10.1016/j.ecoenv.2025.118703

Davis AP et al. 2025. Comparative Toxicogenomics Database’s 20th anniversary: update 2025. Nucleic Acids Research. 53(D1):D1328–D1334. 10.1093/nar/gkae883

Diehl AD et al. 2016. The Cell Ontology 2016: enhanced content, modularization, and ontology interoperability. J Biomed Semant. 7(1):44. 10.1186/s13326-016-0088-7

Farkhondeh T et al. 2020. Organophosphorus Compounds and MAPK Signaling Pathways. IJMS. 21(12):4258. 10.3390/ijms21124258

Feiten AF et al. 2026. TREM2 expression level is critical for microglial state, metabolic capacity and efficacy of TREM2 agonism. Nat Commun. 17(1):2002. 10.1038/s41467-026-68706-8

Gargano MA et al. 2024. The Human Phenotype Ontology in 2024: phenotypes around the world. Nucleic Acids Research. 52(D1):D1333–D1346. 10.1093/nar/gkad1005

Grothe C, Timmer M. 2007. The physiological and pharmacological role of basic fibroblast growth factor in the dopaminergic nigrostriatal system. Brain Research Reviews. 54(1):80–91. 10.1016/j.brainresrev.2006.12.001

Höllerhage M. 2025. Pesticides and parkinson’s disease: causal relationship at the population and individual level? J Neural Transm. [published online ahead of print] [accessed 2026 Jan 14]. https://link.springer.com/10.1007/s00702-025-03054-3. 10.1007/s00702-025-03054-3

Huang H-T, Liao C-K, Chiu W-T, Tzeng S-F. 2017. Ligands of peroxisome proliferator-activated receptor-alpha promote glutamate transporter-1 endocytosis in astrocytes. The International Journal of Biochemistry & Cell Biology. 86:42–53. 10.1016/j.biocel.2017.03.008

Ishii A et al. 2012. ERK1/ERK2 MAPK Signaling is Required to Increase Myelin Thickness Independent of Oligodendrocyte Differentiation and Initiation of Myelination. Journal of Neuroscience. 32(26):8855–8864. 10.1523/JNEUROSCI.0137-12.2012

Ishii A, Furusho M, Dupree JL, Bansal R. 2016. Strength of ERK1/2 MAPK Activation Determines Its Effect on Myelin and Axonal Integrity in the Adult CNS. J Neurosci. 36(24):6471–6487. 10.1523/JNEUROSCI.0299-16.2016

Jaylet T et al. 2025. AOP-helpFinder 3.0: from text mining to network visualization of key event relationships, and knowledge integration from multiple sources Wren J, editor. Bioinformatics. 41(7):btaf381. 10.1093/bioinformatics/btaf381

Karlsson M et al. 2021. A single–cell type transcriptomics map of human tissues. Sci Adv. 7(31):eabh2169. 10.1126/sciadv.abh2169

Kopper TJ, Gensel JC. 2018. Myelin as an inflammatory mediator: Myelin interactions with complement, macrophages, and microglia in spinal cord injury. J Neurosci Res. 96(6):969–977. 10.1002/jnr.24114

Krewski D et al. 2020. Toxicity testing in the 21st century: progress in the past decade and future perspectives. Arch Toxicol. 94(1):1–58. 10.1007/s00204-019-02613-4

López-Merino E, Cuartero MI, Esteban JA, Briz V. 2023. Perinatal exposure to pesticides alters synaptic plasticity signaling and induces behavioral deficits associated with neurodevelopmental disorders. Cell Biol Toxicol. 39(5):2089–2111. 10.1007/s10565-022-09697-2

Luo D et al. 2023. PPARα Inhibits Astrocyte Inflammation Activation by Restoring Autophagic Flux after Transient Brain Ischemia. Biomedicines. 11(3):973. 10.3390/biomedicines11030973

Mak IW, Evaniew N, Ghert M. 2014. Lost in translation: animal models and clinical trials in cancer treatment. Am J Transl Res. 6(2):114–118

Milacic M et al. 2024. The Reactome Pathway Knowledgebase 2024. Nucleic Acids Research. 52(D1):D672–D678. 10.1093/nar/gkad1025

Mortensen HM et al. 2021. The 2021 update of the EPA’s adverse outcome pathway database. Sci Data. 8(1):169. 10.1038/s41597-021-00962-3

Mungall CJ et al. 2012. Uberon, an integrative multi-species anatomy ontology. Genome Biol. 13(1):R5. 10.1186/gb-2012-13-1-r5

OECD. 2017. Revised Guidance Document on Developing and Assessing Adverse Outcome Pathways. Vol 1. OECD Series on Adverse Outcome Pathways Report No.: 184. [accessed 2026 Jan 14]. https://one.oecd.org/document/ENV/JM/MONO(2013)6/en/pdf. 10.1787/5jlv1m9d1g32-en

Oki NO et al. 2016. Accelerating Adverse Outcome Pathway Development Using Publicly Available Data Sources. Curr Envir Health Rpt. 3(1):53–63. 10.1007/s40572-016-0079-y

O’Leary NA et al. 2024. Exploring and retrieving sequence and metadata for species across the tree of life with NCBI Datasets. Sci Data. 11(1):732. 10.1038/s41597-024-03571-y

Ooms J. 2014. The jsonlite Package: A Practical and Consistent Mapping Between JSON Data and R Objects. [accessed 2026 May 29]. https://arxiv.org/abs/1403.2805. 10.48550/ARXIV.1403.2805

Perkins EJ, Woolard EA, Garcia-Reyero N. 2022. Integration of Adverse Outcome Pathways, Causal Networks and ‘Omics to Support Chemical Hazard Assessment. Front Toxicol. 4:786057. 10.3389/ftox.2022.786057

Piñero J et al. 2020. The DisGeNET knowledge platform for disease genomics: 2019 update. Nucleic Acids Research. 48(D1):D845–D855

Putman TE et al. 2024. The Monarch Initiative in 2024: an analytic platform integrating phenotypes, genes and diseases across species. Nucleic Acids Research. 52(D1):D938– D949. 10.1093/nar/gkad1082

R Core Team. 2022. R: A language and environment for statistical computing. R Foundation for Statistical Computing, Vienna, Austria. URL https://www.R-project.org/.

Ramšak Ž et al. 2022. From Causal Networks to Adverse Outcome Pathways: A Developmental Neurotoxicity Case Study. Front Toxicol. 4:815754. 10.3389/ftox.2022.815754

Saarimäki LA et al. 2023. A curated gene and biological system annotation of adverse outcome pathways related to human health. Sci Data. 10(1):409. 10.1038/s41597-023-02321-w

Silge J, Robinson D. 2016. tidytext: Text Mining and Analysis Using Tidy Data Principles in R. JOSS. 1(3):37. 10.21105/joss.00037

Svingen T et al. 2021. A Pragmatic Approach to Adverse Outcome Pathway Development and Evaluation. Toxicological Sciences. 184(2):183–190. 10.1093/toxsci/kfab113

The Gene Ontology Consortium et al. 2023. The Gene Ontology knowledgebase in 2023 Baryshnikova A, editor. GENETICS. 224(1):iyad031. 10.1093/genetics/iyad031

Thomas RS et al. 2019. The Next Generation Blueprint of Computational Toxicology at the U.S. Environmental Protection Agency. Toxicological Sciences. 169(2):317–332. 10.1093/toxsci/kfz058

Timmer M et al. 2007. Fibroblast Growth Factor (FGF)-2 and FGF Receptor 3 Are Required for the Development of the Substantia Nigra, and FGF-2 Plays a Crucial Role for the Rescue of Dopaminergic Neurons after 6-Hydroxydopamine Lesion. J Neurosci. 27(3):459–471. 10.1523/JNEUROSCI.4493-06.2007

Vila M et al. 2001. Bax ablation prevents dopaminergic neurodegeneration in the 1-methyl-4-phenyl-1,2,3,6-tetrahydropyridine mouse model of Parkinson’s disease. Proc Natl Acad Sci USA. 98(5):2837–2842. 10.1073/pnas.051633998

Wang H et al. 2026. FGF–FGFR Signaling in Parkinson’s Disease: Mechanistic Links to Ferroptosis and Neuroprotection. Brain Sciences. 16(2):151. 10.3390/brainsci16020151

Wang Z, Walker GW, Muir DCG, Nagatani-Yoshida K. 2020. Toward a Global Understanding of Chemical Pollution: A First Comprehensive Analysis of National and Regional Chemical Inventories. Environ Sci Technol. 54(5):2575–2584. 10.1021/acs.est.9b06379

Wickham H. 2016. ggplot2: Elegant Graphics for Data Analysis. Springer-Verlag New York. https://ggplot2.tidyverse.org

Wickham H et al. 2019. Welcome to the Tidyverse. JOSS. 4(43):1686. 10.21105/joss.01686

Wickham H. 2023. stringr: Simple, Consistent Wrappers for Common String Operations. https://CRAN.R-project.org/package=stringr

Wickham H et al. dplyr: A Grammar of Data Manipulation_. R package version 1.1.4, <https://CRAN.R-project.org/package=dplyr>.

Wickham H, Pedersen TL, Seidel D. 2023. scales: Scale Functions for Visualization. https://CRAN.R-project.org/package=scales

Wilkins D. 2025. treemapify: Draw Treemaps in ‘ggplot2’. https://CRAN.R-project.org/package=treemapify

Winter DJ. 2017. rentrez: an R package for the NCBI eUtils API. The R Journal. 9(2):520–526

Wu W, Gong X, Qin Z, Wang Y. 2025. Molecular mechanisms of excitotoxicity and their relevance to the pathogenesis of neurodegenerative diseases—an update. Acta Pharmacol Sin. 46(12):3129–3142. 10.1038/s41401-025-01576-w

Yang K et al. 2023. White matter changes in Parkinson’s disease. npj Parkinsons Dis. 9(1):150. 10.1038/s41531-023-00592-z

Yarar N et al. 2024. AOP-networkFinder—a versatile tool for the reconstruction and visualization of Adverse Outcome Pathway networks from AOP-Wiki Fraternali F, editor. Bioinformatics Advances. 5(1):vbaf007. 10.1093/bioadv/vbaf007

Zhang Y, Tan F, Xu P, Qu S. 2016. Recent Advance in the Relationship between Excitatory Amino Acid Transporters and Parkinson’s Disease. Neural Plasticity. 2016:1–8. 10.1155/2016/8941327

