## Supplemental file for "A high-throughput method to computationally develop candidate adverse outcome pathways in humans: a proof of concept with insecticides and Parkinson Disease"

### Supplemental Figures and Tables

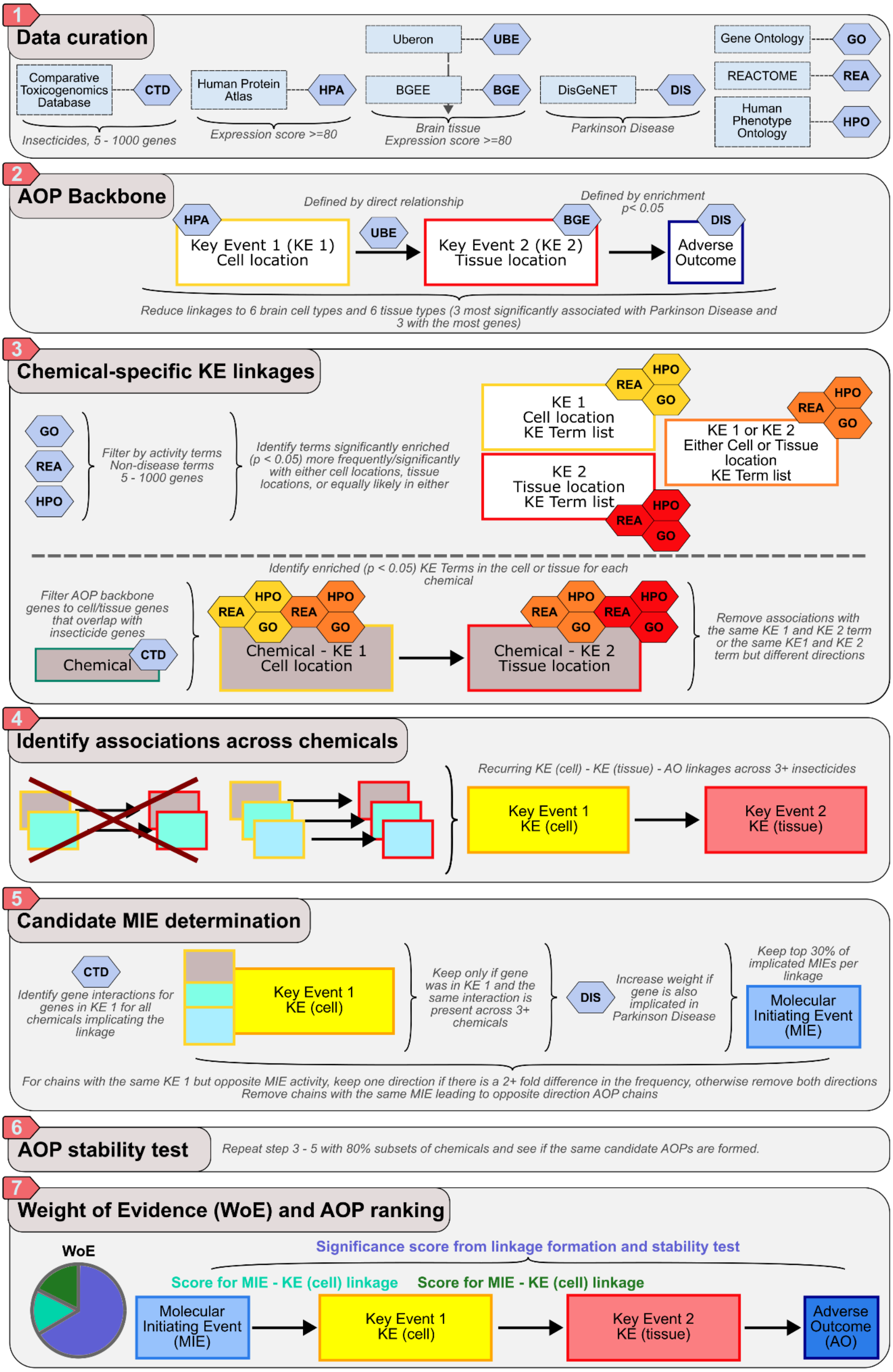


*SI Figure S1. Step by step walkthrough of the AOP construction. Steps 1 – 7 correspond to methods sections 2.1 – 2.7.*

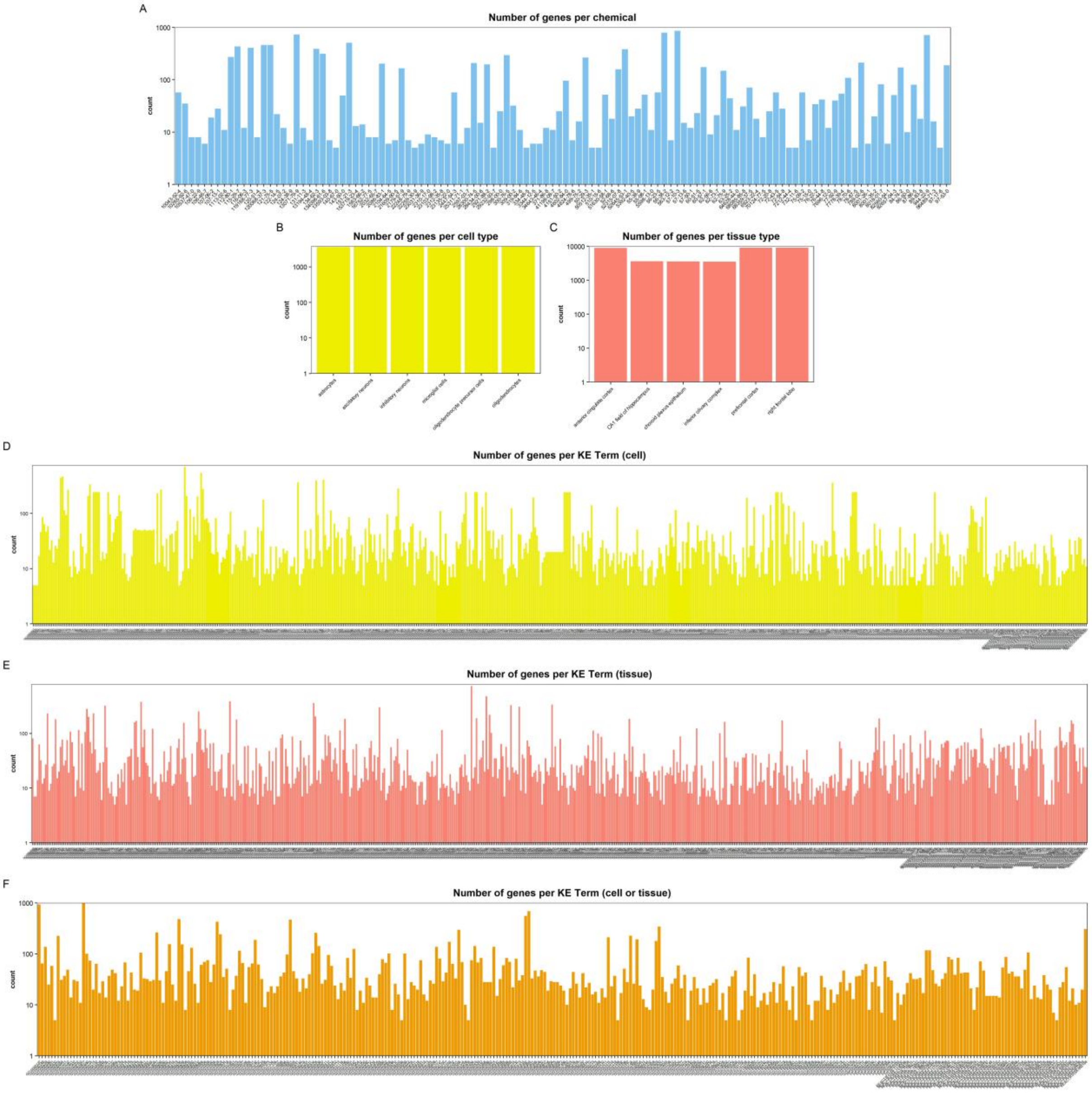


*SI Figure S2. Number of genes per term used to form the linkages. Note that the y-axis is in log10.*

*SI Table S1. Starting databases, versions, and access dates.*

| Resource | Data type | Link | Retrieval Date | Version/Release |
|---|---|---|---|---|
| CTD | Chemical → Gene | https://ctdbase.org/downloads/ | 03/09/2025 | 28/08/2025 |
| GO | Gene ontologies | https://geneontology.org/ | 27/02/2025 | 06/02/2025 |
| REACTOME | Pathways | https://reactome.org/download-data | 29/01/25 | Version 91 |
| HPA | Cell type expression | https://www.proteinatlas.org/humanproteome/single+cell/single+cell+type/data | 22/01/2025 | Version 24.0 |
| MONARCH | Phenotype ontologies | https://monarchinitiative.org/ | 11/02/2026 | February, 2026 |
| NCBI | Human gene set | https://www.ncbi.nlm.nih.gov/datasets/taxonomy/9606/ | 25/02/25 | CLI v16+ (API v2) |
| Uberon | Cell type → Tissues | https://obophenotype.github.io/uberon/current_release/ | 28/03/2025 | / |
| DisGeNET | Gene → Diseases | https://www.disgenet.com/ | 25/06/2025 | Version 25.1 |

*SI Table S2. List of terms used to filter KE Terms by activity that was assembled from the Comparative Toxicogenomics Database chemical – gene interaction action terms.*

| **Activity Terms** |
|---|
| affect |
| react |
| increase |
| express |
| decrease |
| activity |
| chemical synthesi |
| metabolic process |
| transport |
| phosphorylat |
| localizat |
| bind |
| stability |
| response to substance |
| cotreatment |
| abundance |
| hydroxylat |
| glucuronidat |
| cleavage |
| export |
| alkylat |
| secret |
| degradat |
| reduct |
| ubiquitinat |
| oxidat |
| methylat |
| uptake |
| splic |
| mutagenesi |
| fold |
| glutathionylat |
| nitrosat |
| acetylat |
| import |
| sulfat |
| hydrolysi |
| lipidat |
| ADP-ribosylat |
| prenylat |
| ethylat |
| palmitoylat |
| aminat |
| glycosylat |
| farnesylat |
| acylat |
| sumoylat |
| O-linked glycosylat |
| carbamoylat |
| geranoylat |
| glycat |
| carboxylat |
| myristoylat |
| N-linked glycosylat |
| ribosylat |
| polymerizat |
| positive |
| negative |
| regulat |

*SI Table S3. Preliminary literature support for 100 randomly selected candidate AOPs. This support was found by searching for individual KERs with subsequent linkages and locations searched for in the context of pesticide exposure and Parkinson Disease. Ranking for support: 6 = very high (all components of the AOP supported in the literature), 5 = high (only small uncertainties, e.g., support found in the brain but missing explicit tissue location, support found for different nervous system disease than Parkinson Disease), 4 = moderately high (support found for most aspects and the association is plausible, but an explicit link may be missing or feedback loops may be implicated), 3 = moderate (some support, but information may be contradictory or seem to vary depending on different biological contexts), 2 = low (likely not plausible based on preliminary literature, but it is possible it could still apply in certain scenarios), 1 = very low (does not seem plausible). BBB = blood brain barrier, PD = Parkinson Disease, ACC = anterior cingulate cortex, CNS=central nervous system.*

| # | MIETerm | CellTerm | Cell | TissueTerm | Tissue | Score | MIE-KE score | KE-KE score | WoE | Support | Caveats | Overall support |
|---|---|---|---|---|---|---|---|---|---|---|---|---|
| 1 | increases phosphorylation of MAPK8 | positive regulation of gene expression | astrocytes | positive regulation of endothelial cell migration | prefrontal cortex | 0.549 | 0.000 | 1.000 | 0.529 | MAPK8 triggering gene expression in astrocytes, astrocyte dysregulation can affect endothelial cells in the BBB (in prefrontal cortex), e.g., impair tight junction networks[1] and deregulate cell migration, insecticides can do this[2] Pesticides, astrocytes, and BBB implicated in PD[3] | None | **6** |
| 2 | increases phosphorylation of SMAD3 | positive regulation of miRNA transcription | microglial cells | Regulation of PTEN gene transcription | anterior cingulate cortex | 0.574 | 0.000 | 1.000 | 0.544 | SMAD3 regulates PTEN-induced-kinase-1 by positive transactivating transcription, and propose this loop can be targeted in PD interventions[4] SMAD3 regulate microglia[5] | Pathway can be neuroprotective, but insecticides have been seen to alter standard function[6] No direct tissue location support | **3** |
| 3 | increases phosphorylation of MAPK1 | regulation of protein stability | excitatory neurons | regulation of mRNA stability | right frontal lobe | 0.575 | 0.000 | 1.000 | 0.545 | MAPK regulates protein stability[7] and is important in downstream effects in PD[8] and impacts on frontal lobe have been seen from pesticide exposure[3] | No direct cell location support | **5** |
| 4 | increases phosphorylation of EGFR | positive regulation of miRNA transcription | inhibitory neurons | Negative regulation of MAPK pathway | prefrontal cortex | 0.590 | 0.000 | 1.000 | 0.554 | Phosphorylation of EGFR downregulates it[9], and EGFR is downregulated in PD.[10] Dysregulated miRNAs may contribute to PD [11] and negative regulation of MAPK signaling pathway has been seen in the prefrontal cortex with PD-like alterations with depression[12] | No direct cell location support | **5** |
| 5 | increases expression of CTNNB1 | positive regulation of cell population proliferation | microglial cells | regulation of cell cycle | anterior cingulate cortex | 0.529 | 0.000 | 1.000 | 0.517 | β-catenin (coded by CTNNB1) signaling affects microglial activation[13], chronic microglial activation implicated in neurodegeneration[14] | Most discussion around Alzheimer's, No direct tissue location support | **5** |
| 6 | increases phosphorylation of GSK3B | negative regulation of canonical Wnt | excitatory neurons | Regulation of PTEN gene transcription | right frontal lobe | 0.533 | 0.000 | 1.000 | 0.520 | Negative regulation of Wnt via GSK3B by insecticides is implicated in PD[15] and this is observed in excitatory neurons[16] in the cortex in the right frontal lobe and Wnt | Not all pathway support specific to PD | **5** |

| | | | | | | | | | | | | |
|---|---|---|---|---|---|---|---|---|---|---|---|---|
| | | signaling pathway | | | | | | | | signaling is linked to PTEN[17] which is implicated in PD.[18] | | |
| 7 | increases phosphorylation of MAPK1 | protein serine/threonine kinase activity | oligodendrocytes | Detoxification of Reactive Oxygen Species | anterior cingulate cortex | 0.600 | 0.000 | 1.000 | 0.560 | - | Not as clean linkages, and likely describes more positive impacts – a detoxification of ROS would be beneficial to PD | **1** |
| 8 | increases phosphorylation of MAPK1 | protein serine kinase activity | astrocytes | Negative regulation of FGFR3 signaling | anterior cingulate cortex | 0.721 | 0.000 | 1.000 | 0.633 | Organophosphates can sustain MAPK activity[19] and this is upregulated in PD[8] FGFR3 deficient mice struggle to recover from dopaminergic neuron death[20] and dopaminergic neurons are frequent in ACC | FGF signaling can precede MAPK signaling, but MAPK can also feedback on FGF signaling[21] | **4** |
| 9 | decreases expression of ABCA1 | PPARA activates gene expression | astrocytes | positive regulation of cell migration | inferior olivary complex | 0.300 | 0.750 | 1.000 | 0.530 | Decreased ABCA1 can cause astrocyte abnormalities with PPARA pathway implicated[22], ABCA1 deficiency in brain can lead to further damage and is linked to neuroinflammation[23], PPARA is involved in astrocyte autophagic flux[24] and autophagy is a regulator of cell migration[25] | PPARA can also regulate ABCA1, direct tissue link is missing, direct through line not as well-cited as others | **3** |
| 10 | increases expression of NFKB1 | negative regulation of gene expression | inhibitory neurons | Detoxification of Reactive Oxygen Species | right frontal lobe | 0.560 | 0.000 | 1.000 | 0.536 | - | Not as clean linkages, and likely describes more positive impacts – a detoxification of ROS would be beneficial to PD | **1** |
| 11 | increases phosphorylation of MAPK14 | positive regulation of gene expression | oligodendrocytes | RUNX2 regulates osteoblast differentiation | right frontal lobe | 0.529 | 0.000 | 1.000 | 0.517 | - | Highly unlikely with these linkages as osteoblast differentiation relates to bone | **1** |
| 12 | decreases expression of BCL2 | positive regulation of apoptotic process | astrocytes | Regulation of PTEN gene transcription | inferior olivary complex | 0.655 | 0.000 | 1.000 | 0.593 | Pesticides can decrease BCL2 to lead to apoptosis[26] and this decrease in bcl2 can happen in astrocytes[27]. PTEN is implicated in PD.[18] | Connection between BCL2 linked apoptosis and PTEN in the inferior olivary complex is missing – PTEN may precede BCL2 in other cell types[28] | **3** |
| 13 | increases expression of SQSTM1 | positive regulation of apoptotic process | inhibitory neurons | positive regulation of macrophage proliferation | right frontal lobe | 0.569 | 0.000 | 1.000 | 0.541 | Over-expression of SQSTM1 can promote apoptosis via CASP8 in glioma cells,[29] which are in the right frontal lobe.[30] SQSTM1 has been linked to inhibitory neurons[31]. Apoptosis and macrophage proliferation links are implicated in neurodegeneration[32] | All the pieces are there and plausible, but the full connection between them is missing. | **4** |

| | | | | | | | | | | | | |
|---|---|---|---|---|---|---|---|---|---|---|---|---|
| 14 | increases phosphorylation of MAPK8 | positive regulation of gene expression | astrocytes | transcription cis-regulatory region binding | choroid plexus epithelium | 0.511 | 0.000 | 1.000 | 0.507 | MAPK8 triggers gene expression in astrocytes[1] insecticides can cause this[2] and affect BBB while choroid plexus epithelium forms blood-cerebrospinal fluid barrier, which can be affected by inflammation[33] | Fewer citations for direct associations, but pieces of process are supported | **5** |
| 15 | increases phosphorylation of MAPK1 | protein phosphorylation | excitatory neurons | Regulation of PTEN gene transcription | right frontal lobe | 0.774 | 0.000 | 1.000 | 0.665 | MAPK regulates protein phosphorylation in excitatory neurons[34], MAPK pathways can downregulate PTEN[35], PTEN which is implicated in PD.[18,36] | PTEN can also regulate MAPK | **5** |
| 16 | decreases expression of BCL2 | positive regulation of apoptotic process | oligodendrocyte precursor cells | positive regulation of neuroblast proliferation | anterior cingulate cortex | 0.545 | 0.000 | 1.000 | 0.527 | Pesticides can decrease BCL2 to lead to apoptosis[26] | Neuroblast proliferation unlikely in ACC | **2** |
| 17 | increases expression of NFE2L2 | positive regulation of gene expression | astrocytes | Negative regulation of MAPK pathway | CA1 field of hippocampus | 0.604 | 0.000 | 1.000 | 0.563 | Pesticides as triggers NFE2L2 associated w/ pesticides & Parkinson[37], and has been found to help regulate downstream actors in the MAPK pathway (e.g., p38, JNK) in astrocytes[38] which are abundant in the hippocampus | NRF2 can also be regulated by the MAPK pathway | **4** |
| 18 | increases phosphorylation of MAPK8 | positive regulation of gene expression | inhibitory neurons | Negative regulation of FGFR1 signaling | right frontal lobe | 0.601 | 0.000 | 1.000 | 0.560 | Hyperactive RAS/MAPK pathway triggering gene expression in inhibitory neurons[39] and JNK is linked to FGFR in neuron signaling[40], FGFR1 provides neuroprotection and is implicated in PD[41] | While described in neurons, no direct support for MAPK8 pathway in inhibitory neurons, FGF signaling can precede MAPK signaling, but MAPK can also feedback on FGF signaling[21] | **4** |
| 19 | increases phosphorylation of GSK3B | protein serine/threonine kinase activity | microglial cells | Regulation of PTEN gene transcription | CA1 field of hippocampus | 0.592 | 0.000 | 1.000 | 0.555 | Phosphorylation of GSK3B by insecticides is implicated in PD[15] and in microglia-mediated neuroinflammation, and dopamine receptor signaling via Akt.[42] PTEN is linked to the PI3k pathway, is found in the hippocampus,[43] and is implicated in PD[18] | PTEN can also control PI3K/Akt | **4** |
| 20 | increases phosphorylation of MAPK1 | protein serine kinase activity | excitatory neurons | Negative regulation of MAPK pathway | CA1 field of hippocampus | 0.770 | 0.000 | 1.000 | 0.662 | Organophosphates can sustain MAPK activity[19] and this is upregulated in PD[8]. MAPK-mediated protein serine kinase activity is observed in post-mitotic neurons[44] and an increase in MAPK1 leading to negative regulation of the MAPK pathway in the hippocampus has been observed[45] | Exact cell – tissue link missing | **5** |

| | | | | | | | | | | | | |
|---|---|---|---|---|---|---|---|---|---|---|---|---|
| 21 | decreases expression of BCL2 | positive regulation of apoptotic process | excitatory neurons | positive regulation of D-glucose import | inferior olivary complex | 0.572 | 0.000 | 1.000 | 0.543 | Pesticides can decrease BCL2 to lead to apoptosis[26] and the role of BCL2 in excitatory neurons has been studied.[46] Excitotoxic events can lead to glucose import[47] | Exact link between BCL2 and glucose import/apoptosis missing, tissue location support missing | **4** |
| 22 | increases expression of VEGFA | positive regulation of cell population proliferation | oligodendrocyte precursor cells | regulation of cell cycle | prefrontal cortex | 0.524 | 0.000 | 1.000 | 0.514 | VEGFA has been found to promote proliferation and myelination of oligodendrocyte precursor cells[48] and is linked to cell cycle regulation there[49]. Profiles of the prefrontal cortex in PD showed VEGFA pathways upregulated in oligodendrocyte precursor cells[50] | None | **6** |
| 23 | increases expression of JUN | negative regulation of transcription by RNA polymerase II | astrocytes | Regulation of PTEN gene transcription | anterior cingulate cortex | 0.604 | 0.000 | 1.000 | 0.563 | PTEN is implicated in PD.[18] | Increased expression of JUN is more frequently linked to an increase in transcription | **2** |
| 24 | decreases expression of BCL2 | positive regulation of apoptotic process | excitatory neurons | protein serine/threonine/tyrosine kinase activity | prefrontal cortex | 0.564 | 0.000 | 1.000 | 0.539 | Pesticides can decrease BCL2 to lead to apoptosis[26] and the role of BCL2 in excitatory neurons has been studied.[46] BCL2 is linked to protein serine/threonine/tyrosine kinase activity[51,52] and BCL2 mediated neuronal apoptosis has been found in the prefrontal cortex[53] | Can also have protein serine/threonine/tyrosine kinase activity preceding BCL2 | **4** |
| 25 | increases expression of NFKB1 | positive regulation of DNA-templated transcription | oligodendrocytes | Detoxification of Reactive Oxygen Species | inferior olivary complex | 0.560 | 0.000 | 1.000 | 0.536 | - | Not as clean linkages, and likely describes more positive impacts – a detoxification of ROS would be beneficial to PD | **1** |
| 26 | increases expression of JUN | DNA-binding transcription factor activity | microglial cells | transcription cis-regulatory region binding | right frontal lobe | 0.599 | 0.000 | 1.000 | 0.560 | Increased expression of JUN leads to DNA-binding and transcriptional activity and is implicated in neuroinflammation, which is largely driven by microglia.[54], JUN is explicitly in this tissue KE (UniProt). Pesticides (paraquat) affects JUN in microglia[55] | Exact tissue location missing | **5** |
| 27 | increases phosphorylation of MAPK14 | positive regulation of gene expression | inhibitory neurons | RUNX2 regulates osteoblast differentiation | anterior cingulate cortex | 0.537 | 0.000 | 1.000 | 0.522 | - | Highly unlikely with these linkages as osteoblast differentiation relates to bone | **1** |
| 28 | increases expression of VEGFA | positive regulation of blood vessel endothelial cell migration | astrocytes | positive regulation of endothelial cell migration | right frontal lobe | 0.608 | 0.000 | 1.000 | 0.565 | Astrocytes are a primary source for VEGFA in the brain is crucial for new blood vessels and can control migration.[56] This can disrupt BBB integrity ahead of PD[57] | KE term is strongly triggered by this sequence, but does not map directly to astrocytes as opposed | **4** |

| | | | | | | | | | | | | |
|---|---|---|---|---|---|---|---|---|---|---|---|---|
| | | | | | | | | | | | to being triggered by them | |
| 29 | decreases expression of BCL2 | positive regulation of apoptotic process | inhibitory neurons | RNA polymerase II transcription regulator complex | anterior cingulate cortex | 0.580 | 0.000 | 1.000 | 0.548 | Pesticides can decrease BCL2 to lead to apoptosis[26] and the role of BCL2 is implicated in apoptosis of GABAergic neurons[58]. BCL2 linked to RNA Pol II[59] | Tissue link missing | **5** |
| 30 | increases phosphorylation of GSK3B | negative regulation of canonical Wnt signaling pathway | microglial cells | negative regulation of cell growth | CA1 field of hippocampus | 0.510 | 0.000 | 1.000 | 0.506 | Negative regulation of Wnt via GSK3B by insecticides is implicated in PD[15] also involving microglia[60]. WNT signaling components like PP2A involved in cell growth.[61] | Exact tissue location not described, direct WNT – cell growth in brain missing. | **4** |
| 31 | increases phosphorylation of MAPK14 | protein serine/threonine kinase activity | excitatory neurons | protein serine/threonine kinase activator activity | prefrontal cortex | 0.545 | 0.000 | 1.000 | 0.527 | MAPK14 affects excitatory neurons in the cortex and are linked to protein serine/threonine kinase activity, and are implicated in synaptic dysfunction.[62,63] Some serine/threonine activity triggered by MAPK14 (p38) is implicated in PD[64] | Exact tissue location not described | **5** |
| 32 | increases expression of NFKB1 | negative regulation of gene expression | oligodendrocyte precursor cells | Negative feedback regulation of MAPK pathway | CA1 field of hippocampus | 0.512 | 0.000 | 1.000 | 0.507 | Oligodendrocytes are regulated by NFKB1[65] and NFKB1 negatively regulates the MAPK pathway[66] and this pathway is relevant in inflammation in the CA1 field of the hippocampus[67] and in PD[68] | None | **6** |
| 33 | increases expression of NFE2L2 | DNA-binding transcription factor activity | microglial cells | enzyme activator activity | anterior cingulate cortex | 0.590 | 0.000 | 1.000 | 0.554 | Pesticides as triggers NFE2L2 associated w/ pesticides & Parkinson[37], is expressed in microglia and involved in DNA-binding transcription factor activity.[69] Important in enzyme activation[70] | No direct tissue support | **5** |
| 34 | increases expression of PPARG | negative regulation of transcription by RNA polymerase II | microglial cells | negative regulation of miRNA transcription | prefrontal cortex | 0.553 | 0.000 | 1.000 | 0.532 | PPARs can negatively regulate transcription factors, including PPARG in microglial cells.[71] | The pathway is well-described, but tends to be more therapeutic, so adverse pesticide effects likely come from events branching off from this sequence | **3** |
| 35 | increases phosphorylation of MTOR | 3-phosphoinositide-dependent protein kinase activity | excitatory neurons | Negative regulation of MAPK pathway | prefrontal cortex | 0.333 | 0.750 | 1.000 | 0.550 | MTOR can inhibit PI3k pathway[72] and MAPK pathway.[73] These pathways deregulation is important in PD[68]. MTOR is implicated in diverse cell types and neurons, and an increase in phosphorylation of MTOR leads to changes in the prefrontal cortex.[74] | The MTOR – PDK1 process can happen through a feedback loop. | **6** |
| 36 | increases expression of CTNNB1 | positive regulation of apoptotic process | oligodendrocytes | RUNX3 regulates WNT signaling | anterior cingulate cortex | 0.541 | 0.000 | 1.000 | 0.524 | Over expression of B-catenin (encoded by CTNNB1) can lead to apoptosis.[75] The WNT/B-catenin link is important in oliogodendroyctes[76] and regulation by | Not all mechanisms described for references on PD, exact tissue location not described. | **5** |

| | | | | | | | | | | | | |
|---|---|---|---|---|---|---|---|---|---|---|---|---|
| | | | | | | | | | | RUNX3 is linked to CTNNB1 expression[77] and this is important in the brain[78] and PD[79] | | |
| 37 | increases expression of TXNRD1 | PPARA activates gene expression | microglial cells | response to oxidative stress | prefrontal cortex | 0.574 | 0.233 | 1.000 | 0.591 | - | Evidence suggests TXNRD1 negatively regulates PPARA – it is unclear if this would change with sustained pesticide exposure[80] | **2** |
| 38 | increases activity of PPARG | PPARA activates gene expression | inhibitory neurons | protein domain specific binding | choroid plexus epithelium | 0.452 | 0.182 | 1.000 | 0.508 | Pesticides can trigger increases in PPARG and PPARA[81] and these are expressed in inhibitory neurons[82], PPARA triggers gene expression that can use domain-specific binding,[83] and this is relevant in the choroid plexus in nervous system disease.[84] | PPARG does not directly trigger PPARA, so may be more parallel processes | **4** |
| 39 | increases phosphorylation of MAPK1 | regulation of protein stability | oligodendrocytes | Negative regulation of FGFR1 signaling | right frontal lobe | 0.811 | 0.000 | 0.500 | 0.587 | MAPK regulates protein stability[7] and is important in downstream effects in PD[8]. MAPK1 regulates myelin basic protein in oligodendrocytes[85]. FGFR1 signaling linked to MAPK and PD[86] | FGF signaling can precede MAPK signaling, but MAPK can also feedback on FGF signaling[21], no direct tissue description | **4** |
| 40 | increases expression of NFE2L2 | positive regulation of gene expression | astrocytes | RNA polymerase II transcription regulator complex | right frontal lobe | 0.624 | 0.000 | 0.750 | 0.525 | Increased expression of NFE2l2 in astrocytes affects transcription[87] and NRF2 (resulting from NFE2L2) is increased in the brains of PD patients and is important in oxidative stress[88] | Unclear if this process could be dysregulated to be harmful rather than therapeutic. | **3** |
| 41 | increases expression of JUN | positive regulation of miRNA transcription | microglial cells | negative regulation of cell growth | inferior olivary complex | 0.544 | 0.000 | 1.000 | 0.526 | Increased expression of JUN is implicated in neuroinflammation, which is largely driven by microglia[54] and JUN is implicated in increase of miRNAs implicated in Alzheimer's.[89] Upregulated mRNAs following neurotoxicity are implicated in negative regulation of cell growth[90] | Exact links not all described, tissue location not described | **4** |
| 42 | increases expression of NFKB1 | negative regulation of gene expression | microglial cells | Negative regulation of FGFR3 signaling | CA1 field of hippocampus | 0.584 | 0.000 | 1.000 | 0.550 | NFKB1 can negatively regulate gene expression in microglial cells[91] FGFR3 deficient mice struggle to recover from dopaminergic neuron death[20] | The link between NFKB1 and FGFR3 is not clear | **3** |
| 43 | increases expression of NFE2L2 | cellular response to oxidative stress | microglial cells | Negative regulation of FGFR3 signaling | anterior cingulate cortex | 0.573 | 0.000 | 1.000 | 0.544 | NRF2 (from NFE2L2) manages inflammation and response to oxidative stress in microglia.[92] NRF2 is implicated in major signaling pathways[70], and fgfr3 has been mapped to the ACC in lewy body disease[93]. FGFR3 deficient mice struggle to recover from dopaminergic neuron death[20] and dopaminergic neurons are frequent in ACC | Direct link between NFRF2 and FGFR3 is missing, but plausible | **3** |

| | | | | | | | | | | | | |
|---|---|---|---|---|---|---|---|---|---|---|---|---|
| 44 | decreases expression of RB1 | negative regulation of gene expression | astrocytes | regulation of cell cycle | CA1 field of hippocampus | 0.545 | 0.000 | 1.000 | 0.527 | Decreased RB1 expression can lead to further negative regulation of gene expression and RB1 is important in cell cycle, [94] including negative regulation[95] and has been found silenced in astrocytes in gliomas[96]. Aberrant cell cycle in the CA1 field is important in neurodegeneration[97] | Not all evidence directly for PD | **5** |
| 45 | increases phosphorylation of MAPK1 | DNA binding | excitatory neurons | DNA-binding transcription factor binding | anterior cingulate cortex | 0.592 | 0.000 | 1.000 | 0.555 | Environmental toxicants can modulate MAPK1 and excitatory neurons[98] and neurons in the ACC/cortical excitation impacts are important in PD[99] | None | **6** |
| 46 | decreases expression of BCL2 | negative regulation of neuron apoptotic process | astrocytes | Negative regulation of FGFR1 signaling | right frontal lobe | 0.593 | 0.000 | 1.000 | 0.556 | FGFR1 provides neuroprotection and is implicated in PD[41] | Decrease in BCL2 unlikely to lead to negative regulation of apoptotic process | **2** |
| 47 | increases activity of AHR | PPARA activates gene expression | oligodendrocytes | AKT phosphorylates targets in the cytosol | prefrontal cortex | 0.296 | 1.000 | 1.000 | 0.578 | AHR can activate PPARA gene expression[100] and can trigger AKT to phosphorylate targets in cytosol[101], AHRs can disrupt BBB integrity[102] | The AhR – PPARA link is in liver, some question of relevance to oligodendrocytes, some support literature for Alzheimer’s | **3** |
| 48 | increases expression of VIM | positive regulation of gene expression | astrocytes | Negative regulation of FGFR3 signaling | anterior cingulate cortex | 0.600 | 0.000 | 1.000 | 0.560 | Activated astrocytes upregulate filaments like VIM in response to damage.[103] VIM and FGF signaling are linked[104] FGFR3 has been mapped to the ACC in lewy body disease[93]. FGFR3 deficient mice struggle to recover from dopaminergic neuron death[20] and dopaminergic neurons are frequent in ACC | FGFR3 effects may precede VIM | **4** |
| 49 | increases expression of VEGFA | positive regulation of cell population proliferation | astrocytes | positive regulation of endothelial cell chemotaxis | choroid plexus epithelium | 0.623 | 0.000 | 1.000 | 0.574 | VEGFA positively regulates cell population proliferation - astrocyte-derived VEGF stimulates endothelial cell proliferation, but can become deleterious.[105] this could be implicated in the choroid plexus epithelium[106] | Full links across chemotaxis not perfectly clarified. | **5** |
| 50 | increases expression of NFE2L2 | positive regulation of gene expression | microglial cells | Detoxification of Reactive Oxygen Species | right frontal lobe | 0.650 | 0.000 | 1.000 | 0.590 | - | Not as clean linkages, and likely describes more positive impacts – a detoxification of ROS would be beneficial to PD | **1** |
| 51 | decreases expression of BCL2 | positive regulation of apoptotic process | astrocytes | Negative regulation of FGFR1 signaling | CA1 field of hippocampus | 0.573 | 0.000 | 1.000 | 0.544 | Pesticides can decrease BCL2 to lead to apoptosis[26] and this decrease in BCL2 can happen in astrocytes[27]. FGFR1 provides neuroprotection and is | FGF signaling regulates BCL2 normally, but in some cell types/tissues, | **4** |

| | | | | | | | | | | | | |
|---|---|---|---|---|---|---|---|---|---|---|---|---|
| | | | | | | | | | | implicated in PD[41] and this can happen in the hippocampus[107] | there can be feedback loops | |
| 52 | decreases expression of BCL2 | negative regulation of neuron apoptotic process | oligodendrocytes | positive regulation of neuroblast proliferation | prefrontal cortex | 0.507 | 0.000 | 1.000 | 0.504 | - | Decrease in BCL2 unlikely to lead to negative regulation of apoptotic process | **2** |
| 53 | increases phosphorylation of GSK3B | negative regulation of cell migration | astrocytes | positive regulation of endothelial cell migration | prefrontal cortex | 0.517 | 0.000 | 1.000 | 0.510 | GSK3B could regulate cell migration in astrocytes[108] and GSK3B can positively regulate endothelial cell migration[109]. Depending on the phosphorylation site GSK3B can be activated or inactivated and this is relevant in neurodegeneration[110] | A possible parallel process that could happen in the brain that is supported by the behavior of the protein, but unclear if this occurs | **4** |
| 54 | increases expression of VEGFA | positive regulation of angiogenesis | astrocytes | positive regulation of cell migration | choroid plexus epithelium | 0.599 | 0.000 | 1.000 | 0.559 | Astrocyte-mediated angiogenesis with VEGF occurs[111], is implicated in cell migration,[112] | Full throughline to tissue missing | **5** |
| 55 | increases phosphorylation of MAPK8 | positive regulation of gene expression | excitatory neurons | Negative regulation of FGFR3 signaling | right frontal lobe | 0.601 | 0.000 | 1.000 | 0.561 | MAPK8 triggers gene expression[1] insecticides can cause this[2], and the JNK pathway is important for sympathetic neurons.[113] FGFR3 deficient mice struggle to recover from dopaminergic neuron death[20] and MAPK/JNK paths are linked to FGFR3.[114] | Tissue location not confirmed, FGF signaling can precede MAPK signaling, but MAPK can also feedback on FGF signaling[21] | **4** |
| 56 | increases phosphorylation of GSK3B | protein serine kinase activity | microglial cells | Negative regulation of MAPK pathway | prefrontal cortex | 0.578 | 0.000 | 1.000 | 0.547 | Phosphorylation of GSK3B by insecticides is implicated in PD[15] and it is a protein serine kinase present in microglia and implicated in PD.[115] GSK3 is a negative regulator of ERK1/2.[116] Negative regulation of MAPK signaling pathway has been seen in the prefrontal cortex with PD-like alterations with depression[12] | MAPK can modulate GSK3B as well[117], but there is crosstalk | **5** |
| 57 | increases expression of PPARA | PPARA activates gene expression | astrocytes | negative regulation of DNA-templated transcription | prefrontal cortex | 0.250 | 1.000 | 1.000 | 0.550 | PPARA in astrocytes can induce downstream target gene expression[24] and can negatively regulate transcription,[83,118] and astrocytes are common in the prefrontal cortex[119] | None | **6** |
| 58 | increases expression of NFE2L2 | cellular response to oxidative stress | oligodendrocytes | Detoxification of Reactive Oxygen Species | inferior olivary complex | 0.533 | 0.000 | 1.000 | 0.520 | - | Not as clean linkages, and likely describes more positive impacts – a detoxification of ROS would be beneficial to PD | **1** |

| | | | | | | | | | | | | |
|---|---|---|---|---|---|---|---|---|---|---|---|---|
| 59 | decreases expression of BCL2 | positive regulation of apoptotic process | oligodendrocytes | Negative regulation of FGFR1 signaling | CA1 field of hippocampus | 0.668 | 0.000 | 1.000 | 0.601 | Pesticides can decrease BCL2 to lead to apoptosis[26] and this process is relevant in oligodendrocytes.[120] FGFR1 provides neuroprotection and is implicated in PD[41] and this can happen in the hippocampus[107] | The exact BCL2 – FGFR1 link is feasible, but may be in the reverse order | **4** |
| 60 | increases phosphorylation of MAPK14 | negative regulation of canonical Wnt signaling pathway | oligodendrocytes | negative regulation of miRNA transcription | anterior cingulate cortex | 0.512 | 0.000 | 1.000 | 0.507 | MAPK14 (p38alpha) feeds into canonical Wnt via GSK3B[121] and p38 MAPK can regulate Wnt inhibitors in cancer[122] and both pathways are important in oligodendrocytes.[123,124] – negative regulation of Wnt signaling and miRNA changes in neurotoxicity observed with pesticides.[125] | miRNA negative regulation could also modulate Wnt | **5** |
| 61 | increases phosphorylation of SMAD3 | positive regulation of gene expression | oligodendrocyte precursor cells | regulation of cell cycle | anterior cingulate cortex | 0.534 | 0.000 | 1.000 | 0.521 | SMAD3 is targeted by pesticides in the brain and regulates the cell cycle in neurotoxicity.[6] This can happen in oligodendrocyte precursor cells[126] | Tissue linkage missing | **5** |
| 62 | increases activity of PPARG | PPARA activates gene expression | inhibitory neurons | Negative regulation of FGFR1 signaling | CA1 field of hippocampus | 0.597 | 0.182 | 1.000 | 0.595 | Pesticides can trigger increases in PPARG and PPARA[81] and these are expressed in inhibitory neurons[82], FGFR1 provides neuroprotection and is implicated in PD[41] and this can happen in the hippocampus[107] | PPARG does not directly trigger PPARA, while the link between PPARA and FGFR1 is reasonable, it is not explicitly stated. | **3** |
| 63 | affects binding of PPARG | PPARA activates gene expression | inhibitory neurons | protein domain specific binding | CA1 field of hippocampus | 0.573 | 0.000 | 1.000 | 0.544 | Pesticides can trigger increases in PPARG and PPARA[81] and these are expressed in inhibitory neurons[82], PPARA triggers gene expression that can use domain-specific binding,[83] PPARA is expressed in the CA1 field of hippocampus,[127] and this is relevant in nervous system disease.[84] | PPARG does not directly trigger PPARA, may be parallel process | **4** |
| 64 | increases phosphorylation of MAPK14 | positive regulation of gene expression | microglial cells | positive regulation of endothelial cell migration | choroid plexus epithelium | 0.558 | 0.000 | 1.000 | 0.535 | Phosphorylation of p38 increases MMP9 activity which can lead to BBB damage, dysregulation of endothelial tight junctions, and cell migration.[128] The BBB is proximal to the choroid plexus epithelium[129] | Cell and tissue not perfectly clarified | **5** |
| 65 | increases activity of AHR | PPARA activates gene expression | oligodendrocytes | Negative regulation of FGFR1 signaling | CA1 field of hippocampus | 0.235 | 1.000 | 1.000 | 0.541 | Suppression of AHR reduces PPARA in liver.[130] FGFR1 provides neuroprotection and is implicated in PD[41] and this can happen in the hippocampus[107] | This is not the typical dynamic between AHR and PPARA as evidence is limited to liver. Unclear cell influence on this dynamic. while the link between PPARA and FGFR1 is reasonable, it is not explicitly stated. | **3** |

| | | | | | | | | | | | | |
|---|---|---|---|---|---|---|---|---|---|---|---|---|
| 66 | increases expression of SQSTM1 | positive regulation of apoptotic process | oligodendrocytes | RNA polymerase II transcription regulator complex | right frontal lobe | 0.537 | 0.000 | 1.000 | 0.522 | SQSTM1 has a multi-faceted role regulating autophagy and can through this modulate apoptosis[131] and this is important in oligodendrocytes.[132] | The link between sqstm1 and rna pol ii transcription is not unbelievable, but is not documented | **3** |
| 67 | increases phosphorylation of MAPK14 | protein serine kinase activity | excitatory neurons | Negative regulation of FGFR3 signaling | anterior cingulate cortex | 0.530 | 0.000 | 1.000 | 0.518 | P38 MAPK is a serine/threonine protein kinase implicated in modulation of neuronal excitability in neurodegenerative disease.[133] FGFR3 has been mapped to the ACC in lewy body disease[93]. FGFR3 deficient mice struggle to recover from dopaminergic neuron death[20] and dopaminergic neurons are frequent in ACC | FGF signaling can precede MAPK signaling, but MAPK can also feedback on FGF signaling[21] -more clearly connected to FGFR1, but a link to FGFR3 is also feasible | **4** |
| 68 | increases phosphorylation of MAPK8 | positive regulation of gene expression | inhibitory neurons | positive regulation of D-glucose import | right frontal lobe | 0.534 | 0.000 | 1.000 | 0.520 | JNK1 positively regulates gene expression[134] | Connection between JNK1 and glucose import unclear – possible negatively regulates D-glucose import | **2** |
| 69 | increases phosphorylation of MAPK14 | protein serine/threonine kinase activity | astrocytes | Regulation of PTEN gene transcription | prefrontal cortex | 0.539 | 0.000 | 1.000 | 0.524 | P38 MAPK is a serine/threonine protein kinase implicated in neurodegenerative disease in astrocytes[133,135] P38 MAPK is linked to PTEN regulation,[136] and PTEN Important in Parkinson[18], and astrocytes are common and important in the prefrontal cortex[119] | Not all evidence lines directly in brain | **5** |
| 70 | decreases expression of BCL2 | negative regulation of neuron apoptotic process | microglial cells | negative regulation of extrinsic apoptotic signaling pathway | CA1 field of hippocampus | 0.525 | 0.000 | 1.000 | 0.515 | KE link reasonable | Decrease in BCL2 unlikely to lead to negative regulation of apoptotic process | **2** |
| 71 | increases phosphorylation of MAPK1 | protein phosphorylation | oligodendrocytes | Regulation of PTEN gene transcription | choroid plexus epithelium | 0.669 | 0.000 | 1.000 | 0.601 | Organophosphates can sustain MAPK activity[19], this pathway is important in PD[137] and MAPK1 is important in oligodendrocytes[138], MAPK pathways can downregulate PTEN[35], PTEN Important in Important in Parkinson[18] | Tissue location missing, PTEN can also regulate MAPK | **5** |
| 72 | increases expression of NFKB1 | positive regulation of DNA-templated transcription | astrocytes | TP53 Regulates Metabolic Genes | anterior cingulate cortex | 0.560 | 0.000 | 1.000 | 0.536 | insecticides trigger NFKB to promote oxidative stress[139], NFKB1 is a transcription factor important to astrocytes.[140] TP53 is triggered via NFKB signaling and can increase oxidative stress in the brain[141] | Direct tissue location not confirmed | **5** |

| | | | | | | | | | | | | |
|---|---|---|---|---|---|---|---|---|---|---|---|---|
| 73 | increases expression of FAS | positive regulation of apoptotic process | astrocytes | Negative regulation of FGFR3 signaling | CA1 field of hippocampus | 0.546 | 0.000 | 1.000 | 0.528 | FAS mediates apoptosis and is in astrocytes.[142] FGFR3 deficient mice struggle to recover from dopaminergic neuron death[20] | Human astrocytes may be protected against FAS-induced apoptosis without a mechanism to bypass it[143]. The link between FAS and FGFR3 is plausible but needs further assessment | **4** |
| 74 | increases expression of NFKB1 | negative regulation of transcription by RNA polymerase II | microglial cells | RNA polymerase II transcription regulator complex | CA1 field of hippocampus | 0.563 | 0.000 | 1.000 | 0.538 | Homodimers of the p50 subunit of NFKB can inhibit transcription[144] and NFKB-p50 subunits are found in microglia[91]. NFKB transcription crucial to numerous diseases | The exact dynamics between each step need to be further clarified, but appear plausible | **4** |
| 75 | increases phosphorylation of GSK3B | protein serine kinase activity | oligodendrocyte precursor cells | Regulation of PTEN gene transcription | anterior cingulate cortex | 0.599 | 0.000 | 1.000 | 0.559 | Phosphorylation of GSK3B by insecticides is implicated in PD[15] and it is a protein serine kinase present in oligodendrocyte precursor cells[145] and regulates PTEN gene transcription.[146] PTEN Important in Parkinson[18] | Tissue location not named | **5** |
| 76 | increases phosphorylation of GSK3B | RNA polymerase II-specific DNA-binding transcription factor binding | excitatory neurons | DNA-binding transcription factor binding | prefrontal cortex | 0.513 | 0.000 | 1.000 | 0.508 | GSK3B regulates many transcription factors via RNA polymerase II activity[147,148] and is important in hippocampal neurons[149] | Exact cell and tissue support absent | **5** |
| 77 | increases expression of FOS | positive regulation of miRNA transcription | astrocytes | RUNX2 regulates osteoblast differentiation | CA1 field of hippocampus | 0.530 | 0.000 | 1.000 | 0.518 | - | Highly unlikely with these linkages as osteoblast differentiation relates to bone | **1** |
| 78 | increases phosphorylation of GSK3B | protein serine/threonine kinase activity | excitatory neurons | protein serine/threonine kinase activator activity | anterior cingulate cortex | 0.523 | 0.000 | 1.000 | 0.514 | Phosphorylation of GSK3B by insecticides is implicated in PD[15] and it is a protein serine kinase present in excitatory neurons and implicated in PD.[115] GSK3B is in the ACC[150] | None | **6** |

| | | | | | | | | | | | | |
|---|---|---|---|---|---|---|---|---|---|---|---|---|
| 79 | increases phosphorylation of SMAD3 | RNA polymerase II-specific DNA-binding transcription factor binding | oligodendrocyte precursor cells | DNA-binding transcription factor binding | right frontal lobe | 0.543 | 0.000 | 1.000 | 0.526 | SMAD3 is important in oligodendrocyte precursor cells[126] and is a DNA-binding transcription factor.[151,152] | Direct link to PD currently missing – other unnamed components likely additionally involved | **4** |
| 80 | increases expression of SQSTM1 | positive regulation of apoptotic process | microglial cells | circadian regulation of gene expression | choroid plexus epithelium | 0.538 | 0.000 | 1.000 | 0.523 | SQSTM1 has a multi-faceted role regulating autophagy and can through this modulate apoptosis and are in microglia[153]. BMAL1 regulates circadian gene expression, and SQSTM1 interacts with ARNTL/BMAL1 in the brain and may be important in neurodegenerative disease.[154,155] | Exact tissue location missing | **5** |
| 81 | increases phosphorylation of MAPK1 | DNA binding | astrocytes | enzyme binding | inferior olivary complex | 0.683 | 0.000 | 0.501 | 0.510 | Organophosphates can sustain MAPK activity[19] and this is upregulated in PD[8]. Phosphorylation of ERK1/2 influences transcription in astrocytes[156] and MAPK1 is important in enzyme binding[157] | Tissue support missing | **5** |
| 82 | increases phosphorylation of MAPK1 | protein serine/threonine kinase activity | excitatory neurons | Negative regulation of FGFR3 signaling | anterior cingulate cortex | 0.746 | 0.000 | 1.000 | 0.647 | Organophosphates can sustain MAPK activity[19] and this is upregulated in PD[8]. MAPK-mediated protein serine kinase activity is observed in post-mitotic neurons[44]. fgfr3 has been mapped to the ACC in lewy body disease[93]. FGFR3 deficient mice struggle to recover from dopaminergic neuron death[20] and dopaminergic neurons are frequent in ACC | FGF signaling can precede MAPK signaling, but MAPK can also feedback on FGF signaling[21] | **4** |
| 83 | increases expression of PPARA | PPARA activates gene expression | astrocytes | MAP kinase activity | anterior cingulate cortex | 0.541 | 1.000 | 1.000 | 0.724 | PPARA is a transcription factor that regulates many processes[158] and is implicated in astrocyte behavior in Alzheimer's.[159]<br>PPARA is linked to MAPK activity[160] and MAPK signaling is implicated in PD[161] | Cross-talk between pathways, tissue support missing | **5** |
| 84 | increases phosphorylation of MAPK1 | regulation of protein stability | oligodendrocytes | Regulation of PTEN gene transcription | prefrontal cortex | 0.797 | 0.000 | 1.000 | 0.678 | Organophosphates can sustain MAPK activity[19], MAPK signaling regulates protein stability[7] and is important in downstream effects in PD[8]<br>MAPK1 is important in oligodendrocytes[138], MAPK pathways can downregulate PTEN[35],<br>PTEN Important in Parkinson[18] and prefrontal cortex[162] | PTEN can also regulate MAPK | **5** |

| | | | | | | | | | | | | |
|---|---|---|---|---|---|---|---|---|---|---|---|---|
| 85 | increases phosphorylation of MAPK8 | positive regulation of apoptotic process | oligodendrocyte precursor cells | RNA polymerase II transcription regulator complex | inferior olivary complex | 0.509 | 0.000 | 1.000 | 0.505 | MAPK/JNK link to apoptosis[36] and is important in oligodendrocyte precursor cells[163] and in transcription regulation[164] | Exact tissue location missing | **5** |
| 86 | increases phosphorylation of MAPK1 | protein phosphorylation | astrocytes | positive regulation of protein autophosphorylation | prefrontal cortex | 0.577 | 0.000 | 1.000 | 0.546 | Organophosphates can sustain MAPK activity[19], this pathway is important in PD and astrocytes[137] and this is upregulated in PD[8], and can induce autophosphorylation[134] and astrocytes are common in the prefrontal cortex[119] | None | **6** |
| 87 | increases phosphorylation of GSK3B | negative regulation of canonical Wnt signaling pathway | astrocytes | Negative feedback regulation of MAPK pathway | prefrontal cortex | 0.599 | 0.000 | 1.000 | 0.559 | Negative regulation of Wnt via GSK3B by insecticides is implicated in PD[15], this is linked to MAPK signaling,[165] regulation of MAPK signaling pathway has been seen in the prefrontal cortex with PD-like alterations with depression[12], and astrocytes are common in the prefrontal cortex[119] | None | **6** |
| 88 | increases phosphorylation of GSK3B | negative regulation of cell migration | oligodendrocyte precursor cells | positive regulation of macrophage proliferation | prefrontal cortex | 0.586 | 0.000 | 1.000 | 0.552 | GSK3B regulates cell migration[108] and models have simulated GSK3B knockout triggering macrophage proliferation[166] | Directionality of GSK3B depends on where its phosphorylated – this mechanism is plausible but requires more assessment, cell location and tissue location unclear | **3** |
| 89 | increases expression of PPARA | positive regulation of DNA-templated transcription | astrocytes | Negative regulation of FGFR3 signaling | CA1 field of hippocampus | 0.533 | 0.000 | 1.000 | 0.520 | PPARA is a transcription factor that regulates many processes[158] and is implicated in astrocyte behavior in Alzheimer's.[159] FGFR3 deficient mice struggle to recover from dopaminergic neuron death[20] | The link between FGFR3 and PPARA is plausible but not outlined. | **4** |
| 90 | increases phosphorylation of MAPK8 | protein phosphorylation | excitatory neurons | Regulation of PTEN gene transcription | choroid plexus epithelium | 0.541 | 0.000 | 1.000 | 0.525 | MAPK/JNK link phosphorylation in excitatory neurons[167] and regulates PTEN[168] which can upregulate neuron death [169], important in Parkinson[18] | choroid plexus epithelium support missing | **5** |
| 91 | increases phosphorylation of MAPK14 | negative regulation of canonical Wnt signaling pathway | inhibitory neurons | negative regulation of miRNA transcription | anterior cingulate cortex | 0.505 | 0.000 | 1.000 | 0.503 | MAPK14 (p38alpha) feeds into canonical Wnt via GSK3B[121] and p38 MAPK can regulate Wnt inhibitors in cancer[122] – negative regulation of Wnt signaling and miRNA changes in neurotoxicity observed with pesticides.[125] | miRNA negative regulation could also modulate Wnt, cell evidence less clear | **4** |
| 92 | increases expression of HIF1A | positive regulation of | astrocytes | response to oxidative stress | choroid plexus epithelium | 0.589 | 0.000 | 1.000 | 0.553 | HIF1A can activate gene transcription.[170] HIF1A was expressed at high levels in | Most evidence relates to protection, but dysregulation from | **3** |

| | | | | | | | | | | | | |
|---|---|---|---|---|---|---|---|---|---|---|---|---|
| | | DNA-templated transcription | | | | | | | | the choroid plexus of mice[171] and is linked to oxidative stress response.[172] | pesticides occurs in other systems[173] | |
| 93 | increases expression of FOS | positive regulation of miRNA transcription | microglial cells | transcription cis-regulatory region binding | choroid plexus epithelium | 0.547 | 0.000 | 1.000 | 0.528 | FOS positively regulates miRNA signaling[174], including miRNAs important in microglia, transcription regulation, and PD.[175] | Tissue confirmation not clear | **5** |
| 94 | increases phosphorylation of MTOR | protein serine kinase activity | oligodendrocytes | Negative regulation of FGFR1 signaling | CA1 field of hippocampus | 0.506 | 0.000 | 1.000 | 0.504 | MTOR possesses serine-threonine protein kinase activity and is essential to coordinate myelination in oligodendrocytes.[176] mTOR regulates FGF21 which mediates its effects by stimulating FGFR1.[177] FGFR1 provides neuroprotection and is implicated in PD[41] and this can happen in the hippocampus[107] | MTOR – FGF linkage not in brain, may involve negative feedback or behave differently in different regions. | **3** |
| 95 | increases phosphorylation of MAPK14 | positive regulation of erythrocyte differentiation | microglial cells | Negative regulation of MAPK pathway | CA1 field of hippocampus | 0.530 | 0.000 | 1.000 | 0.518 | Activation of p38 MAPK induces erythroid differentiation and inhibition of ERK[178] negative regulation of the MAPK pathway in the hippocampus has been observed[45] | Negative regulation of MAPK pathway may happen in parallel, the cell location may not align | **3** |
| 96 | increases phosphorylation of MAPK14 | positive regulation of gene expression | excitatory neurons | positive regulation of D-glucose import | choroid plexus epithelium | 0.557 | 0.000 | 1.000 | 0.534 | MAPK14 can mediate positively regulate gene expression and glycolytic flux.[179] this gene expression regulation can happen in neurons[180] | Tissue location not directly evidenced | **5** |
| 97 | increases phosphorylation of MAPK1 | peptidyl-serine phosphorylation | astrocytes | protein serine/threonine/tyrosine kinase activity | right frontal lobe | 0.574 | 0.000 | 1.000 | 0.545 | Organophosphates can sustain MAPK activity[19], this pathway is important in PD and astrocytes[137] and this is upregulated in PD[8], MAPK pathway modulates protein serine/threonine/tyrosine kinase activity[181] | Not a perfect tissue link | **5** |
| 98 | increases expression of NFE2L2 | positive regulation of gene expression | astrocytes | positive regulation of D-glucose import | inferior olivary complex | 0.559 | 0.000 | 1.000 | 0.535 | Pesticides as triggers NFE2L2 associated w/ pesticides & Parkinson[37], NFE2L2 triggering expression in astrocytes and regulation of glucose uptake[182] | Direct link to tissue location absent | **5** |
| 99 | increases phosphorylation of MAPK8 | positive regulation of apoptotic process | excitatory neurons | Regulation of PTEN gene transcription | choroid plexus epithelium | 0.542 | 0.000 | 1.000 | 0.525 | MAPK/JNK link to apoptosis and PTEN[36], regulates PTEN[168] important in neuron death[169] Important in Parkinson[18] | Direct link to tissue location absent, | **5** |
| 100 | decreases expression of SOD1 | positive regulation of apoptotic process | oligodendrocytes | RNA polymerase II transcription regulator complex | inferior olivary complex | 0.537 | 0.000 | 1.000 | 0.522 | SOD1 decrease increases oxidative stress and triggers apoptosis[183], Disrupted SOD1 triggers oligodendrocyte apoptosis and motor neuron degeneration[184]Decreased SOD1 from pesticides and Parkinson[185] | Direct link to tissue location absent, GxE interactions may be important | **5** |

**Case study references**


1. Wu D, Ma Y, Wang B, et al. Pathological Roles of Astrocytes in Traumatic Brain Injury. *CNS Neurosci Ther*. 2026;32(4):e70856. doi:10.1002/cns.70856
2. Maurya SK, Mishra J, Tripathi VK, Sharma R, Siddiqui MH. Cypermethrin induces astrocyte damage: Role of aberrant Ca2+, ROS, JNK, P38, matrix metalloproteinase 2 and migration related reelin protein. *Pestic Biochem Physiol*. 2014;111:51-59. doi:10.1016/j.pestbp.2014.03.005
3. Sharma N, An SSA. Soil to Synapse: Molecular Insights into the Neurotoxicity of Common Gardening Chemicals in Alzheimer's and Parkinson's Disease. *Int J Mol Sci*. 2025;26(13):6468. doi:10.3390/ijms26136468
4. Tang M, Rong D, Gao X, et al. A positive feedback loop between SMAD3 and PINK1 in regulation of mitophagy. *Cell Discov*. 2025;11(1):22. doi:10.1038/s41421-025-00774-4
5. Liu Y, Yu L, Xu Y, Tang X, Wang X. Substantia nigra Smad3 signaling deficiency: relevance to aging and Parkinson's disease and roles of microglia, proinflammatory factors, and MAPK. *J Neuroinflammation*. 2020;17(1):342. doi:10.1186/s12974-020-02023-9
6. Seth B, Yadav A, Agarwal S, Tiwari SK, Chaturvedi RK. Inhibition of the transforming growth factor-β/SMAD cascade mitigates the anti-neurogenic effects of the carbamate pesticide carbofuran. *J Biol Chem*. 2017;292(47):19423-19440. doi:10.1074/jbc.M117.798074
7. Vargas-Ibarra D, Velez-Vasquez M, Bermudez-Munoz M. Regulation of MAPK ERK1/2 Signaling by Phosphorylation: Implications in Physiological and Pathological Contexts. In: Ying S, ed. *Post-Translational Modifications in Cellular Functions and Diseases*. IntechOpen; 2021. doi:10.5772/intechopen.97061
8. Dzamko N, Zhou J, Huang Y, Halliday GM. Parkinsonâ€™s disease-implicated kinases in the brain; insights into disease pathogenesis. *Front Mol Neurosci*. 2014;7. doi:10.3389/fnmol.2014.00057
9. Ahsan A, Hiniker SM, Ramanand SG, et al. Role of Epidermal Growth Factor Receptor Degradation in Cisplatin-Induced Cytotoxicity in Head and Neck Cancer. *Cancer Res*. 2010;70(7):2862-2869. doi:10.1158/0008-5472.CAN-09-4294
10. Mencke P, Hanss Z, Boussaad I, Sugier PE, Elbaz A, Krüger R. Bidirectional Relation Between Parkinson's Disease and Glioblastoma Multiforme. *Front Neurol*. 2020;11:898. doi:10.3389/fneur.2020.00898
11. Nies YH, Mohamad Najib NH, Lim WL, Kamaruzzaman MA, Yahaya MF, Teoh SL. MicroRNA Dysregulation in Parkinson's Disease: A Narrative Review. *Front Neurosci*. 2021;15:660379. doi:10.3389/fnins.2021.660379
12. Ma Z, Xu Y, Lian P, et al. Alpha-synuclein Fibrils Inhibit Activation of the BDNF/ERK Signaling Loop in the mPFC to Induce Parkinson's Disease-like Alterations with Depression. *Neurosci Bull*. 2025;41(6):951-969. doi:10.1007/s12264-024-01323-x
13. Zheng H, Jia L, Liu CC, et al. TREM2 Promotes Microglial Survival by Activating Wnt/β-Catenin Pathway. *J Neurosci Off J Soc Neurosci*. 2017;37(7):1772-1784. doi:10.1523/JNEUROSCI.2459-16.2017
14. Smith JA, Das A, Ray SK, Banik NL. Role of pro-inflammatory cytokines released from microglia in neurodegenerative diseases. *Brain Res Bull*. 2012;87(1):10-20. doi:10.1016/j.brainresbull.2011.10.004
15. Zhang L, Cen L, Qu S, et al. Enhancing Beta-Catenin Activity via GSK3beta Inhibition Protects PC12 Cells against Rotenone Toxicity through Nurr1 Induction. Le W, ed. *PLOS ONE*. 2016;11(4):e0152931. doi:10.1371/journal.pone.0152931
16. Marosi M, Arman P, Aceto G, D'Ascenzo M, Laezza F. Glycogen Synthase Kinase 3: Ion Channels, Plasticity, and Diseases. *Int J Mol Sci*. 2022;23(8):4413. doi:10.3390/ijms23084413
17. Persad A, Venkateswaran G, Hao L, et al. Active β-catenin is regulated by the PTEN/PI3 kinase pathway: a role for protein phosphatase PP2A. *Genes Cancer*. 2017;7(11-12):368-382. doi:10.18632/genesandcancer.128
18. Yang X, Liu T, Cheng H. PTEN: a new dawn in Parkinson's disease treatment. *Front Cell Neurosci*. 2025;19:1497555. doi:10.3389/fncel.2025.1497555

19. Farkhondeh T, Mehrpour O, Buhrmann C, Pourbagher-Shahri AM, Shakibaei M, Samarghandian S. Organophosphorus Compounds and MAPK Signaling Pathways. *Int J Mol Sci*. 2020;21(12):4258. doi:10.3390/ijms21124258
20. Timmer M, Cesnulevicius K, Winkler C, et al. Fibroblast Growth Factor (FGF)-2 and FGF Receptor 3 Are Required for the Development of the Substantia Nigra, and FGF-2 Plays a Crucial Role for the Rescue of Dopaminergic Neurons after 6-Hydroxydopamine Lesion. *J Neurosci*. 2007;27(3):459-471. doi:10.1523/JNEUROSCI.4493-06.2007
21. Zakrzewska M, Haugsten EM, Nadratowska-Wesolowska B, et al. ERK-Mediated Phosphorylation of Fibroblast Growth Factor Receptor 1 on Ser$^{777}$ Inhibits Signaling. *Sci Signal*. 2013;6(262). doi:10.1126/scisignal.2003087
22. Shinozaki Y, Leung A, Namekata K, et al. Astrocytic dysfunction induced by ABCA1 deficiency causes optic neuropathy. *Sci Adv*. 2022;8(44):eabq1081. doi:10.1126/sciadv.abq1081
23. Cui X, Chopp M, Zacharek A, et al. Deficiency of brain ATP-binding cassette transporter A-1 exacerbates blood-brain barrier and white matter damage after stroke. *Stroke*. 2015;46(3):827-834. doi:10.1161/STROKEAHA.114.007145
24. Luo D, Ye W, Chen L, et al. PPARα Inhibits Astrocyte Inflammation Activation by Restoring Autophagic Flux after Transient Brain Ischemia. *Biomedicines*. 2023;11(3):973. doi:10.3390/biomedicines11030973
25. Kenific CM, Wittmann T, Debnath J. Autophagy in adhesion and migration. *J Cell Sci*. 2016;129(20):3685-3693. doi:10.1242/jcs.188490
26. Jang YJ, Won JH, Back MJ, et al. Paraquat Induces Apoptosis through a Mitochondria-Dependent Pathway in RAW264.7 Cells. *Biomol Ther*. 2015;23(5):407-413. doi:10.4062/biomolther.2015.075
27. Xu M, Zhang H ling. Death and survival of neuronal and astrocytic cells in ischemic brain injury: a role of autophagy. *Acta Pharmacol Sin*. 2011;32(9):1089-1099. doi:10.1038/aps.2011.50
28. Huang H, Cheville JC, Pan Y, Roche PC, Schmidt LJ, Tindall DJ. PTEN Induces Chemosensitivity in PTEN-mutated Prostate Cancer Cells by Suppression of Bcl-2 Expression. *J Biol Chem*. 2001;276(42):38830-38836. doi:10.1074/jbc.M103632200
29. Zhang YB, Gong JL, Xing TY, Zheng SP, Ding W. Autophagy protein p62/SQSTM1 is involved in HAMLET-induced cell death by modulating apotosis in U87MG cells. *Cell Death Dis*. 2013;4(3):e550-e550. doi:10.1038/cddis.2013.77
30. Larjavaara S, Mäntylä R, Salminen T, et al. Incidence of gliomas by anatomic location. *Neuro-Oncol*. 2007;9(3):319-325. doi:10.1215/15228517-2007-016
31. Hui KK, Tanaka M. Autophagy links MTOR and GABA signaling in the brain. *Autophagy*. 2019;15(10):1848-1849. doi:10.1080/15548627.2019.1637643
32. Cui J, Zhao S, Li Y, et al. Regulated cell death: discovery, features and implications for neurodegenerative diseases. *Cell Commun Signal*. 2021;19(1):120. doi:10.1186/s12964-021-00799-8
33. Liu R, Zhang Z, Chen Y, et al. Choroid plexus epithelium and its role in neurological diseases. *Front Mol Neurosci*. 2022;15:949231. doi:10.3389/fnmol.2022.949231
34. Yuan LL, Adams JP, Swank M, Sweatt JD, Johnston D. Protein Kinase Modulation of Dendritic $K^+$ Channels in Hippocampus Involves a Mitogen-Activated Protein Kinase Pathway. *J Neurosci*. 2002;22(12):4860-4868. doi:10.1523/JNEUROSCI.22-12-04860.2002
35. Salmena L, Carracedo A, Pandolfi PP. Tenets of PTEN Tumor Suppression. *Cell*. 2008;133(3):403-414. doi:10.1016/j.cell.2008.04.013
36. Walker CL, Liu NK, Xu XM. PTEN/PI3K and MAPK signaling in protection and pathology following CNS injuries. *Front Biol*. 2013;8(4):421-433. doi:10.1007/s11515-013-1255-1
37. Paul KC, Sinsheimer JS, Cockburn M, Bronstein JM, Bordelon Y, Ritz B. NFE2L2, PPARGC1α, and pesticides and Parkinson's disease risk and progression. *Mech Ageing Dev*. 2018;173:1-8. doi:10.1016/j.mad.2018.04.004

38. Lin YT, Wu PH, Tsai YC, et al. Indoxyl Sulfate Induces Apoptosis Through Oxidative Stress and Mitogen-Activated Protein Kinase Signaling Pathway Inhibition in Human Astrocytes. *J Clin Med*. 2019;8(2):191. doi:10.3390/jcm8020191
39. Knowles SJ, Stafford AM, Zaman T, et al. Distinct hyperactive RAS/MAPK alleles converge on common GABAergic interneuron core programs. *Development*. 2023;150(10):dev201371. doi:10.1242/dev.201371
40. Castro-Torres RD, Olloquequi J, Parcerisas A, et al. JNK signaling and its impact on neural cell maturation and differentiation. *Life Sci*. 2024;350:122750. doi:10.1016/j.lfs.2024.122750
41. Wang J, Xing L, Song Y, et al. FGFs/FGFRs and neuroinflammatory diseases: mechanism, drug therapies and delivery systems. *J Adv Res*. 2025;78:733-756. doi:10.1016/j.jare.2025.06.024
42. Yu H, Xiong M, Zhang Z. The role of glycogen synthase kinase 3 beta in neurodegenerative diseases. *Front Mol Neurosci*. 2023;16:1209703. doi:10.3389/fnmol.2023.1209703
43. Wang Y, Cheng A, Mattson MP. The PTEN Phosphatase is Essential for Long-Term Depression of Hippocampal Synapses. *NeuroMolecular Med*. 2006;8(3):329-336. doi:10.1385/NMM:8:3:329
44. Mao LM, Wang JQ. Regulation of Group I Metabotropic Glutamate Receptors by MAPK/ERK in Neurons. *J Nat Sci*. 2016;2(12):e268.
45. Geng M, Shao Q, Fu J, et al. Down-regulation of MKP-1 in hippocampus protects against stress-induced depression-like behaviors and neuroinflammation. *Transl Psychiatry*. 2024;14(1):130. doi:10.1038/s41398-024-02846-7
46. Schelman WR, Andres RD, Sipe KJ, Kang E, Weyhenmeyer JA. Glutamate mediates cell death and increases the Bax to Bcl-2 ratio in a differentiated neuronal cell line. *Mol Brain Res*. 2004;128(2):160-169. doi:10.1016/j.molbrainres.2004.06.011
47. Weisová P, Concannon CG, Devocelle M, Prehn JHM, Ward MW. Regulation of glucose transporter 3 surface expression by the AMP-activated protein kinase mediates tolerance to glutamate excitation in neurons. *J Neurosci Off J Soc Neurosci*. 2009;29(9):2997-3008. doi:10.1523/JNEUROSCI.0354-09.2009
48. Li R, Wang H, Sun N, et al. VEGF-A promotes the proliferation and myelination of induced oligodendrocyte progenitor cells in rat spinal cord injury lesions. *Brain Res Bull*. 2025;231:111554. doi:10.1016/j.brainresbull.2025.111554
49. Choi EH, Xu Y, Medynets M, et al. Activated T cells induce proliferation of oligodendrocyte progenitor cells via release of vascular endothelial cell growth factor-A. *Glia*. 2018;66(11):2503-2513. doi:10.1002/glia.23501
50. Zhu B, Park JM, Coffey SR, et al. Single-cell transcriptomic and proteomic analysis of Parkinson’s disease brains. *Sci Transl Med*. 2024;16(771):eabo1997. doi:10.1126/scitranslmed.abo1997
51. Gajewski TF, Thompson CB. Apoptosis Meets Signal Transduction: Elimination of a BAD Influence. *Cell*. 1996;87(4):589-592. doi:10.1016/S0092-8674(00)81377-X
52. Cui J, Placzek WJ. Post-Transcriptional Regulation of Anti-Apoptotic BCL2 Family Members. *Int J Mol Sci*. 2018;19(1):308. doi:10.3390/ijms19010308
53. Li Y, Han F, Shi Y. Increased Neuronal Apoptosis in Medial Prefrontal Cortex is Accompanied with Changes of Bcl-2 and Bax in a Rat Model of Post-Traumatic Stress Disorder. *J Mol Neurosci*. 2013;51(1):127-137. doi:10.1007/s12031-013-9965-z
54. Khan FA, Khan H, Awan UA, et al. c-Jun in neurodegeneration: A key transcriptional regulator with therapeutic implications. *Mol Ther Nucleic Acids*. 2026;37(2):102874. doi:10.1016/j.omtn.2026.102874
55. Sun Y, Zheng J, Xu Y, Zhang X. Paraquat-induced inflammatory response of microglia through HSP60/TLR4 signaling. *Hum Exp Toxicol*. 2018;37(11):1161-1168. doi:10.1177/0960327118758152
56. Bozoyan L, Khlghatyan J, Saghatelyan A. Astrocytes Control the Development of the Migration-Promoting Vasculature Scaffold in the Postnatal Brain via VEGF Signaling. *J Neurosci*. 2012;32(5):1687-1704. doi:10.1523/JNEUROSCI.5531-11.2012

57. Cabezas R, Ã□vila M, Gonzalez J, et al. Astrocytic modulation of blood brain barrier: perspectives on Parkinsonâ€™s disease. *Front Cell Neurosci*. 2014;8. doi:10.3389/fncel.2014.00211
58. Wong FK, Bercsenyi K, Sreenivasan V, Portalés A, Fernández-Otero M, Marín O. Pyramidal cell regulation of interneuron survival sculpts cortical networks. *Nature*. 2018;557(7707):668-673. doi:10.1038/s41586-018-0139-6
59. Harper NW, Birdsall GA, Honeywell ME, Ward KM, Pai AA, Lee MJ. RNA Pol II inhibition activates cell death independently from the loss of transcription. *Cell*. 2025;188(22):6301-6316.e29. doi:10.1016/j.cell.2025.07.034
60. L'Episcopo F, Tirolo C, Serapide MF, et al. Microglia Polarization, Gene-Environment Interactions and Wnt/β-Catenin Signaling: Emerging Roles of Glia-Neuron and Glia-Stem/Neuroprogenitor Crosstalk for Dopaminergic Neurorestoration in Aged Parkinsonian Brain. *Front Aging Neurosci*. 2018;10:12. doi:10.3389/fnagi.2018.00012
61. Qin K, Yu M, Fan J, et al. Canonical and noncanonical Wnt signaling: Multilayered mediators, signaling mechanisms and major signaling crosstalk. *Genes Dis*. 2024;11(1):103-134. doi:10.1016/j.gendis.2023.01.030
62. Falcicchia C, Tozzi F, Arancio O, Watterson DM, Origlia N. Involvement of p38 MAPK in Synaptic Function and Dysfunction. *Int J Mol Sci*. 2020;21(16):5624. doi:10.3390/ijms21165624
63. Mao LM, Wang JQ. Synaptically Localized Mitogen-Activated Protein Kinases: Local Substrates and Regulation. *Mol Neurobiol*. 2016;53(9):6309-6315. doi:10.1007/s12035-015-9535-1
64. Chen J, Ren Y, Gui C, et al. Phosphorylation of Parkin at serine 131 by p38 MAPK promotes mitochondrial dysfunction and neuronal death in mutant A53T α-synuclein model of Parkinson's disease. *Cell Death Dis*. 2018;9(6):700. doi:10.1038/s41419-018-0722-7
65. Blank T, Prinz M. NF-ÎºB signaling regulates myelination in the CNS. *Front Mol Neurosci*. 2014;7. doi:10.3389/fnmol.2014.00047
66. Beinke S, Deka J, Lang V, et al. NF-kappaB1 p105 negatively regulates TPL-2 MEK kinase activity. *Mol Cell Biol*. 2003;23(14):4739-4752. doi:10.1128/MCB.23.14.4739-4752.2003
67. Galvis-Montes DS, van Loo KMJ, van Waardenberg AJ, et al. Highly dynamic inflammatory and excitability transcriptional profiles in hippocampal CA1 following status epilepticus. *Sci Rep*. 2023;13(1):22187. doi:10.1038/s41598-023-49310-y
68. Jha SK, Jha NK, Kar R, Ambasta RK, Kumar P. p38 MAPK and PI3K/AKT Signalling Cascades inParkinson's Disease. *Int J Mol Cell Med*. 2015;4(2):67-86.
69. De Plano LM, Calabrese G, Rizzo MG, Oddo S, Caccamo A. The Role of the Transcription Factor Nrf2 in Alzheimer's Disease: Therapeutic Opportunities. *Biomolecules*. 2023;13(3):549. doi:10.3390/biom13030549
70. He F, Ru X, Wen T. NRF2, a Transcription Factor for Stress Response and Beyond. *Int J Mol Sci*. 2020;21(13):4777. doi:10.3390/ijms21134777
71. Ricote M, Glass CK. PPARs and molecular mechanisms of transrepression. *Biochim Biophys Acta*. 2007;1771(8):926-935. doi:10.1016/j.bbalip.2007.02.013
72. Dibble CC, Cantley LC. Regulation of mTORC1 by PI3K signaling. *Trends Cell Biol*. 2015;25(9):545-555. doi:10.1016/j.tcb.2015.06.002
73. Efeyan A, Sabatini DM. mTOR and cancer: many loops in one pathway. *Curr Opin Cell Biol*. 2010;22(2):169-176. doi:10.1016/j.ceb.2009.10.007
74. Lipton JO, Sahin M. The Neurology of mTOR. *Neuron*. 2014;84(2):275-291. doi:10.1016/j.neuron.2014.09.034
75. Kim K, Pang KM, Evans M, Hay ED. Overexpression of beta-catenin induces apoptosis independent of its transactivation function with LEF-1 or the involvement of major G1 cell cycle regulators. *Mol Biol Cell*. 2000;11(10):3509-3523. doi:10.1091/mbc.11.10.3509
76. Vallée A, Vallée JN, Guillevin R, Lecarpentier Y. Interactions Between the Canonical WNT/Beta-Catenin Pathway and PPAR Gamma on Neuroinflammation, Demyelination,

and Remyelination in Multiple Sclerosis. *Cell Mol Neurobiol*. 2018;38(4):783-795. doi:10.1007/s10571-017-0550-9
77. Ito K. RUNX3 in oncogenic and anti-oncogenic signaling in gastrointestinal cancers. *J Cell Biochem*. 2011;112(5):1243-1249. doi:10.1002/jcb.23047
78. Zhuang W, Ye T, Wang W, Song W, Tan T. CTNNB1 in neurodevelopmental disorders. *Front Psychiatry*. 2023;14:1143328. doi:10.3389/fpsyt.2023.1143328
79. Serafino A, Cozzolino M. The Wnt/β-catenin signaling: a multifunctional target for neuroprotective and regenerative strategies in Parkinson’s disease. *Neural Regen Res*. 2023;18(2):306-308. doi:10.4103/1673-5374.343908
80. Liu GH, Qu J, Shen X. Thioredoxin-mediated Negative Autoregulation of Peroxisome Proliferator-activated Receptor α Transcriptional Activity. *Mol Biol Cell*. 2006;17(4):1822-1833. doi:10.1091/mbc.e05-10-0979
81. Hernández-Valdez J, Velázquez-Zepeda A, Sánchez-Meza JC. Effect of Pesticides on Peroxisome Proliferator-Activated Receptors (PPARs) and Their Association with Obesity and Diabetes. *PPAR Res*. 2023;2023:1743289. doi:10.1155/2023/1743289
82. Xi ZX, Hempel B, Crissman M, et al. PPARα and PPARγ are expressed in midbrain dopamine neurons and modulate dopamine- and cannabinoid-mediated behavior in mice. Preprint posted online March 2, 2023:rs. 3.rs-2614714. doi:10.21203/rs.3.rs-2614714/v1
83. Galanou ON, Konstandi M. Unraveling the Function of PPARα in Neurodegenerative Disorders: A Potential Pathway to Novel Therapies. *Biomedicines*. 2025;13(11):2813. doi:10.3390/biomedicines13112813
84. Stopa EG, Tanis KQ, Miller MC, et al. Comparative transcriptomics of choroid plexus in Alzheimer’s disease, frontotemporal dementia and Huntington’s disease: implications for CSF homeostasis. *Fluids Barriers CNS*. 2018;15(1):18. doi:10.1186/s12987-018-0102-9
85. Ishii A, Furusho M, Dupree JL, Bansal R. Strength of ERK1/2 MAPK Activation Determines Its Effect on Myelin and Axonal Integrity in the Adult CNS. *J Neurosci*. 2016;36(24):6471-6487. doi:10.1523/JNEUROSCI.0299-16.2016
86. Wang H, Wen X, Yan M, Li R, Mao D, Tian X. FGF–FGFR Signaling in Parkinson’s Disease: Mechanistic Links to Ferroptosis and Neuroprotection. *Brain Sci*. 2026;16(2):151. doi:10.3390/brainsci16020151
87. Calkins MJ, Vargas MR, Johnson DA, Johnson JA. Astrocyte-Specific Overexpression of Nrf2 Protects Striatal Neurons from Mitochondrial Complex II Inhibition. *Toxicol Sci*. 2010;115(2):557-568. doi:10.1093/toxsci/kfq072
88. Petri S, Körner S, Kiaei M. Nrf2/ARE Signaling Pathway: Key Mediator in Oxidative Stress and Potential Therapeutic Target in ALS. *Neurol Res Int*. 2012;2012:1-7. doi:10.1155/2012/878030
89. Guedes JR, Custódia CM, Silva RJ, De Almeida LP, Pedroso De Lima MC, Cardoso AL. Early miR-155 upregulation contributes to neuroinflammation in Alzheimer’s disease triple transgenic mouse model. *Hum Mol Genet*. 2014;23(23):6286-6301. doi:10.1093/hmg/ddu348
90. Sarkar S, Pandey A, Khan B, Pant AB. Integrative assessment of RNA sequencing and in silico analysis to pinpoint mRNAs, lncRNAs, and circRNAs interactions with miRNAs underlying arsenic-induced neurotoxicity. *BMC Genomics*. 2025;26(1):813. doi:10.1186/s12864-025-11970-7
91. Taetzsch T, Levesque S, McGraw C, et al. Redox regulation of NF-κB p50 and M1 polarization in microglia. *Glia*. 2015;63(3):423-440. doi:10.1002/glia.22762
92. He X, Dando O, Qiu J. Nrf2 controls homeostatic transcriptional signatures and inflammatory responses in a cell-type specific manner in the adult mouse brain. *iScience*. 2025;28(9):113198. doi:10.1016/j.isci.2025.113198
93. Tsuchida T, Mano T, Koshi-Mano K, et al. Methylation changes and aberrant expression of FGFR3 in Lewy body disease neurons. *Brain Res*. 2018;1697:59-66. doi:10.1016/j.brainres.2018.06.017

94. Janostiak R, Torres-Sanchez A, Posas F, de Nadal E. Understanding Retinoblastoma Post-Translational Regulation for the Design of Targeted Cancer Therapies. *Cancers*. 2022;14(5):1265. doi:10.3390/cancers14051265
95. Indovina P, Pentimalli F, Casini N, Vocca I, Giordano A. RB1 dual role in proliferation and apoptosis: cell fate control and implications for cancer therapy. *Oncotarget*. 2015;6(20):17873-17890. doi:10.18632/oncotarget.4286
96. Pavlou MAS, Grandbarbe L, Buckley NJ, Niclou SP, Michelucci A. Transcriptional and epigenetic mechanisms underlying astrocyte identity. *Prog Neurobiol*. 2019;174:36-52. doi:10.1016/j.pneurobio.2018.12.007
97. Koseoglu MM, Norambuena A, Sharlow ER, Lazo JS, Bloom GS. Aberrant Neuronal Cell Cycle Re-Entry: The Pathological Confluence of Alzheimer's Disease and Brain Insulin Resistance, and Its Relation to Cancer. *J Alzheimers Dis JAD*. 2019;67(1):1-11. doi:10.3233/JAD-180874
98. Ijomone OM, Iroegbu JD, Aschner M, Bornhorst J. Impact of environmental toxicants on p38- and ERK-MAPK signaling pathways in the central nervous system. *Neurotoxicology*. 2021;86:166-171. doi:10.1016/j.neuro.2021.08.005
99. Zhou Z, Ye P, Li XH, et al. Synaptic potentiation of anterior cingulate cortex contributes to chronic pain of Parkinson's disease. *Mol Brain*. 2021;14(1):161. doi:10.1186/s13041-021-00870-y
100. Oshida K, Vasani N, Thomas RS, et al. Screening a mouse liver gene expression compendium identifies modulators of the aryl hydrocarbon receptor (AhR). *Toxicology*. 2015;336:99-112. doi:10.1016/j.tox.2015.07.005
101. Coumoul X, Barouki R, Esser C, et al. The aryl hydrocarbon receptor: structure, signaling, physiology and pathology. *Signal Transduct Target Ther*. 2026;11(1):20. doi:10.1038/s41392-025-02500-8
102. Al-kuraishy HM, Al-Maiahy TJ, Sulaiman GM, et al. The potential role of aryl hydrocarbon receptor in Alzheimer's disease: Protective or detrimental. *Ageing Res Rev*. 2025;110:102810. doi:10.1016/j.arr.2025.102810
103. Kang W, Hébert JM. Signaling Pathways in Reactive Astrocytes, a Genetic Perspective. *Mol Neurobiol*. 2011;43(3):147-154. doi:10.1007/s12035-011-8163-7
104. Kang W, Balordi F, Su N, Chen L, Fishell G, Hébert JM. Astrocyte activation is suppressed in both normal and injured brain by FGF signaling. *Proc Natl Acad Sci*. 2014;111(29). doi:10.1073/pnas.1320401111
105. Egervari K, Potter G, Guzman-Hernandez ML, et al. Astrocytes spatially restrict VEGF signaling by polarized secretion and incorporation of VEGF into the actively assembling extracellular matrix. *Glia*. 2016;64(3):440-456. doi:10.1002/glia.22939
106. Solár P, Zamani A, Kubíčková L, Dubový P, Joukal M. Choroid plexus and the blood-cerebrospinal fluid barrier in disease. *Fluids Barriers CNS*. 2020;17(1):35. doi:10.1186/s12987-020-00196-2
107. Shin J, Oh H, Park JH, et al. Dysregulation of FGFR1 signaling in the hippocampus facilitates depressive disorder. *Exp Mol Med*. 2025;57(8):1818-1836. doi:10.1038/s12276-025-01519-9
108. Sun T, Rodriguez M, Kim L. Glycogen synthase kinase 3 in the world of cell migration. *Dev Growth Differ*. 2009;51(9):735-742. doi:10.1111/j.1440-169X.2009.01141.x
109. Santio NM, Salmela M, Arola H, et al. The PIM1 kinase promotes prostate cancer cell migration and adhesion via multiple signalling pathways. *Exp Cell Res*. 2016;342(2):113-124. doi:10.1016/j.yexcr.2016.02.018
110. Krishnankutty A, Kimura T, Saito T, et al. In vivo regulation of glycogen synthase kinase 3β activity in neurons and brains. *Sci Rep*. 2017;7(1):8602. doi:10.1038/s41598-017-09239-5
111. He Z, Mao Y, Chi L, Sun J. Astrocyte-mediated angiogenesis in CNS diseases: Mechanisms and therapeutic implications. *Brain Res Bull*. 2026;240:111914. doi:10.1016/j.brainresbull.2026.111914
112. Chapouly C, Tadesse Argaw A, Horng S, et al. Astrocytic TYMP and VEGFA drive blood-brain barrier opening in inflammatory central nervous system lesions. *Brain J Neurol*. 2015;138(Pt 6):1548-1567. doi:10.1093/brain/awv077

113. Kristiansen M, Hughes R, Patel P, Jacques TS, Clark AR, Ham J. Mkp1 is a c-Jun target gene that antagonizes JNK-dependent apoptosis in sympathetic neurons. *J Neurosci Off J Soc Neurosci*. 2010;30(32):10820-10832. doi:10.1523/JNEUROSCI.2824-10.2010
114. Ornitz DM, Itoh N. The Fibroblast Growth Factor signaling pathway. *Wiley Interdiscip Rev Dev Biol*. 2015;4(3):215-266. doi:10.1002/wdev.176
115. Lai S, Wang P, Gong J, Zhang S. New insights into the role of GSK-3β in the brain: from neurodegenerative disease to tumorigenesis. *PeerJ*. 2023;11:e16635. doi:10.7717/peerj.16635
116. Wang Q, Zhou Y, Wang X, Evers BM. Glycogen synthase kinase-3 is a negative regulator of extracellular signal-regulated kinase. *Oncogene*. 2006;25(1):43-50. doi:10.1038/sj.onc.1209004
117. Thornton TM, Pedraza-Alva G, Deng B, et al. Phosphorylation by p38 MAPK as an Alternative Pathway for GSK3β Inactivation. *Science*. 2008;320(5876):667-670. doi:10.1126/science.1156037
118. Yang Y, Gocke AR, Lovett-Racke A, Drew PD, Racke MK. PPAR Alpha Regulation of the Immune Response and Autoimmune Encephalomyelitis. *PPAR Res*. 2008;2008:546753. doi:10.1155/2008/546753
119. Kim E, Du H, Tan Y, et al. Prefrontal cortex astrocytes modulate distinct neuronal populations to control anxiety-like behavior. *Nat Commun*. 2025;16(1):7819. doi:10.1038/s41467-025-63131-9
120. Butts BD, Houde C, Mehmet H. Maturation-dependent sensitivity of oligodendrocyte lineage cells to apoptosis: implications for normal development and disease. *Cell Death Differ*. 2008;15(7):1178-1186. doi:10.1038/cdd.2008.70
121. Lepore Signorile M, Fasano C, Forte G, et al. Uncoupling p38α nuclear and cytoplasmic functions and identification of two p38α phosphorylation sites on β-catenin: implications for the Wnt signaling pathway in CRC models. *Cell Biosci*. 2023;13(1):223. doi:10.1186/s13578-023-01175-4
122. Browne AJ, Göbel A, Thiele S, Hofbauer LC, Rauner M, Rachner TD. p38 MAPK regulates the Wnt inhibitor Dickkopf-1 in osteotropic prostate cancer cells. *Cell Death Dis*. 2016;7(2):e2119-e2119. doi:10.1038/cddis.2016.32
123. Dai ZM, Sun S, Wang C, et al. Stage-Specific Regulation of Oligodendrocyte Development by Wnt/ -Catenin Signaling. *J Neurosci*. 2014;34(25):8467-8473. doi:10.1523/JNEUROSCI.0311-14.2014
124. Chew LJ, Coley W, Cheng Y, Gallo V. Mechanisms of Regulation of Oligodendrocyte Development by p38 Mitogen-Activated Protein Kinase. *J Neurosci*. 2010;30(33):11011-11027. doi:10.1523/JNEUROSCI.2546-10.2010
125. Yu G, Su Q, Chen Y, Wu L, Wu S, Li H. Epigenetics in neurodegenerative disorders induced by pesticides. *Genes Environ Off J Jpn Environ Mutagen Soc*. 2021;43(1):55. doi:10.1186/s41021-021-00224-z
126. Palazuelos J, Klingener M, Aguirre A. TGF Signaling Regulates the Timing of CNS Myelination by Modulating Oligodendrocyte Progenitor Cell Cycle Exit through SMAD3/4/FoxO1/Sp1. *J Neurosci*. 2014;34(23):7917-7930. doi:10.1523/JNEUROSCI.0363-14.2014
127. Roy A, Jana M, Corbett GT, et al. Regulation of cyclic AMP response element binding and hippocampal plasticity-related genes by peroxisome proliferator-activated receptor α. *Cell Rep*. 2013;4(4):724-737. doi:10.1016/j.celrep.2013.07.028
128. Lao D, Gong Z, Li T, Mo X, Huang W. The P38MAPK Pathway Mediates the Destruction of the Blood–Brain Barrier in Anti-NMDAR Encephalitis Mice. *Neurochem Res*. 2025;50(1):21. doi:10.1007/s11064-024-04270-1
129. Engelhardt B, Sorokin L. The blood–brain and the blood–cerebrospinal fluid barriers: function and dysfunction. *Semin Immunopathol*. 2009;31(4):497-511. doi:10.1007/s00281-009-0177-0
130. Wang C, Xu CX, Krager SL, Bottum KM, Liao DF, Tischkau SA. Aryl Hydrocarbon Receptor Deficiency Enhances Insulin Sensitivity and Reduces PPAR-α Pathway

Activity in Mice. *Environ Health Perspect*. 2011;119(12):1739-1744. doi:10.1289/ehp.1103593
131. Hou Y, Liu L, Zhao T, Guo Y, Shi J. The multifaceted role of SQSTM1/p62 in disc degeneration: A master regulator of cellular stress responses. *Biochem Biophys Rep*. 2025;44:102222. doi:10.1016/j.bbrep.2025.102222
132. Zhang T, Bhambri A, Zhang Y, et al. Autophagy collaborates with apoptosis pathways to control oligodendrocyte number. *Cell Rep*. 2023;42(8):112943. doi:10.1016/j.celrep.2023.112943
133. Lee JK, Kim NJ. Recent Advances in the Inhibition of p38 MAPK as a Potential Strategy for the Treatment of Alzheimer's Disease. *Molecules*. 2017;22(8):1287. doi:10.3390/molecules22081287
134. Cargnello M, Roux PP. Activation and function of the MAPKs and their substrates, the MAPK-activated protein kinases. *Microbiol Mol Biol Rev MMBR*. 2011;75(1):50-83. doi:10.1128/MMBR.00031-10
135. Bachstetter AD, Van Eldik LJ. The p38 MAP Kinase Family as Regulators of Proinflammatory Cytokine Production in Degenerative Diseases of the CNS. *Aging Dis*. 2010;1(3):199-211.
136. Kim YC, Gonzalez-Nieves R, Cutler ML. Rsu1-dependent control of PTEN expression is regulated via ATF2 and cJun. *J Cell Commun Signal*. 2019;13(3):331-341. doi:10.1007/s12079-018-00504-4
137. Kim EK, Choi EJ. Pathological roles of MAPK signaling pathways in human diseases. *Biochim Biophys Acta BBA - Mol Basis Dis*. 2010;1802(4):396-405. doi:10.1016/j.bbadis.2009.12.009
138. Bhat NR, Zhang P. Activation of Mitogen-Activated Protein Kinases in Oligodendrocytes. *J Neurochem*. 1996;66(5):1986-1994. doi:10.1046/j.1471-4159.1996.66051986.x
139. Gargouri B, Boukholda K, Kumar A, et al. Bifenthrin insecticide promotes oxidative stress and increases inflammatory mediators in human neuroblastoma cells through NF-kappaB pathway. *Toxicol In Vitro*. 2020;65:104792. doi:10.1016/j.tiv.2020.104792
140. Ageeva T, Rizvanov A, Mukhamedshina Y. NF-κB and JAK/STAT Signaling Pathways as Crucial Regulators of Neuroinflammation and Astrocyte Modulation in Spinal Cord Injury. *Cells*. 2024;13(7):581. doi:10.3390/cells13070581
141. Zhang K, Zhao Y, Chen X, et al. p53 promote oxidative stress, neuroinflammation and behavioral disorders via DDIT4-NF-κB signaling pathway. *Redox Biol*. 2025;86:103836. doi:10.1016/j.redox.2025.103836
142. Saas P, Boucraut J, Quiquerez AL, et al. CD95 (Fas/Apo-1) as a Receptor Governing Astrocyte Apoptotic or Inflammatory Responses: A Key Role in Brain Inflammation? *J Immunol*. 1999;162(4):2326-2333. doi:10.4049/jimmunol.162.4.2326
143. Song JH, Bellail A, Tse MCL, Yong VW, Hao C. Human astrocytes are resistant to Fas ligand and tumor necrosis factor-related apoptosis-inducing ligand-induced apoptosis. *J Neurosci Off J Soc Neurosci*. 2006;26(12):3299-3308. doi:10.1523/JNEUROSCI.5572-05.2006
144. Elsharkawy AM, Oakley F, Lin F, Packham G, Mann DA, Mann J. The NF-kappaB p50:p50:HDAC-1 repressor complex orchestrates transcriptional inhibition of multiple pro-inflammatory genes. *J Hepatol*. 2010;53(3):519-527. doi:10.1016/j.jhep.2010.03.025
145. Zhou L, Shao CY, Xu S min, et al. GSK3β Promotes the Differentiation of Oligodendrocyte Precursor Cells via β-Catenin-Mediated Transcriptional Regulation. *Mol Neurobiol*. 2014;50(2):507-519. doi:10.1007/s12035-014-8678-9
146. Bermúdez Brito M. Focus on PTEN regulation. *Front Oncol*. 2015;5. doi:10.3389/fonc.2015.00166
147. Nieto Moreno N, Villafañez F, Giono LE, et al. GSK-3 is an RNA polymerase II phospho-CTD kinase. *Nucleic Acids Res*. 2020;48(11):6068-6080. doi:10.1093/nar/gkaa322
148. Beurel E, Grieco SF, Jope RS. Glycogen synthase kinase-3 (GSK3): regulation, actions, and diseases. *Pharmacol Ther*. 2015;148:114-131. doi:10.1016/j.pharmthera.2014.11.016

149. Engel T, Gómez-Sintes R, Alves M, et al. Bi-directional genetic modulation of GSK-3β exacerbates hippocampal neuropathology in experimental status epilepticus. *Cell Death Dis*. 2018;9(10):969. doi:10.1038/s41419-018-0963-5
150. Wang M, Liu X, Hou Y, et al. Decrease of GSK-3β Activity in the Anterior Cingulate Cortex of Shank3b−/− Mice Contributes to Synaptic and Social Deficiency. *Front Cell Neurosci*. 2019;13:447. doi:10.3389/fncel.2019.00447
151. Dickey TH, Pyle AM. The SMAD3 transcription factor binds complex RNA structures with high affinity. *Nucleic Acids Res*. 2017;45(20):11980-11988. doi:10.1093/nar/gkx846
152. Inman GJ. Linking Smads and transcriptional activation. *Biochem J*. 2005;386(Pt 1):e1-e3. doi:10.1042/bj20042133
153. Zhang R, Kang R, Tang D. SQSTM1/p62 at the Crossroads of Autophagy, Inflammation, and Lethal Infection. *Cells*. 2026;15(7):652. doi:10.3390/cells15070652
154. Ding Z, Zhuang Y, Jian J, et al. The role of BMAL1 in glial cells: Implications for cognitive function in neurodegenerative diseases. *Brain Res Bull*. 2025;229:111463. doi:10.1016/j.brainresbull.2025.111463
155. Yang M, Chen P, Liu J, et al. Clockophagy is a novel selective autophagy process favoring ferroptosis. *Sci Adv*. 2019;5(7):eaaw2238. doi:10.1126/sciadv.aaw2238
156. Krueger J, Chou FL, Glading A, Schaefer E, Ginsberg MH. Phosphorylation of Phosphoprotein Enriched in Astrocytes (PEA-15) Regulates Extracellular Signal-regulated Kinase-dependent Transcription and Cell Proliferation. *Mol Biol Cell*. 2005;16(8):3552-3561. doi:10.1091/mbc.e04-11-1007
157. Motta M, Pannone L, Pantaleoni F, et al. Enhanced MAPK1 Function Causes a Neurodevelopmental Disorder within the RASopathy Clinical Spectrum. *Am J Hum Genet*. 2020;107(3):499-513. doi:10.1016/j.ajhg.2020.06.018
158. Rakhshandehroo M, Knoch B, Müller M, Kersten S. Peroxisome Proliferator-Activated Receptor Alpha Target Genes. *PPAR Res*. 2010;2010:1-20. doi:10.1155/2010/612089
159. Raha S, Ghosh A, Dutta D, Patel DR, Pahan K. Activation of PPARα enhances astroglial uptake and degradation of β-amyloid. *Sci Signal*. 2021;14(706):eabg4747. doi:10.1126/scisignal.abg4747
160. Meng R, Pei Z, Zhang A, et al. AMPK activation enhances PPARα activity to inhibit cardiac hypertrophy via ERK1/2 MAPK signaling pathway. *Arch Biochem Biophys*. 2011;511(1-2):1-7. doi:10.1016/j.abb.2011.04.010
161. Bohush A, Niewiadomska G, Filipek A. Role of Mitogen Activated Protein Kinase Signaling in Parkinson’s Disease. *Int J Mol Sci*. 2018;19(10):2973. doi:10.3390/ijms19102973
162. Wang XQ, Zhang L, Xia ZY, Chen JY, Fang Y, Ding YQ. PTEN in prefrontal cortex is essential in regulating depression-like behaviors in mice. *Transl Psychiatry*. 2021;11(1):185. doi:10.1038/s41398-021-01312-y
163. Lorenzati M, Boda E, Parolisi R, et al. c-Jun N-terminal kinase 1 (JNK1) modulates oligodendrocyte progenitor cell architecture, proliferation and myelination. *Sci Rep*. 2021;11(1):7264. doi:10.1038/s41598-021-86673-6
164. Yue J, López JM. Understanding MAPK Signaling Pathways in Apoptosis. *Int J Mol Sci*. 2020;21(7):2346. doi:10.3390/ijms21072346
165. Xue C, Chu Q, Shi Q, Zeng Y, Lu J, Li L. Wnt signaling pathways in biology and disease: mechanisms and therapeutic advances. *Signal Transduct Target Ther*. 2025;10(1):106. doi:10.1038/s41392-025-02142-w
166. Zerrouk N, Alcraft R, Hall BA, Augé F, Niarakis A. Large-scale computational modelling of the M1 and M2 synovial macrophages in rheumatoid arthritis. *Npj Syst Biol Appl*. 2024;10(1):10. doi:10.1038/s41540-024-00337-5
167. Hong Y, Siino V, Flinkman D, et al. JNK-regulated phosphoproteome links synaptic and metabolic pathways to mood regulation. *Neurobiol Dis*. 2026;218:107207. doi:10.1016/j.nbd.2025.107207
168. Hettinger K, Vikhanskaya F, Poh MK, et al. c-Jun promotes cellular survival by suppression of PTEN. *Cell Death Differ*. 2007;14(2):218-229. doi:10.1038/sj.cdd.4401946

169. Kim AH, Yano H, Cho H, et al. Akt1 Regulates a JNK Scaffold during Excitotoxic Apoptosis. *Neuron*. 2002;35(4):697-709. doi:10.1016/S0896-6273(02)00821-8
170. Cimmino F, Avitabile M, Lasorsa VA, et al. HIF-1 transcription activity: HIF1A driven response in normoxia and in hypoxia. *BMC Med Genet*. 2019;20(1):37. doi:10.1186/s12881-019-0767-1
171. Kratzer I, Liddelow SA, Saunders NR, Dziegielewska KM, Strazielle N, Ghersi-Egea JF. Developmental changes in the transcriptome of the rat choroid plexus in relation to neuroprotection. *Fluids Barriers CNS*. 2013;10(1):25. doi:10.1186/2045-8118-10-25
172. Li HS, Zhou YN, Li L, et al. HIF-1α protects against oxidative stress by directly targeting mitochondria. *Redox Biol*. 2019;25:101109. doi:10.1016/j.redox.2019.101109
173. Xiao S, Cui J, Cao Y, et al. Adolescent exposure to organophosphate insecticide malathion induces spermatogenesis dysfunction in mice by activating the HIF-1/MAPK/PI3K pathway. *Environ Pollut*. 2024;363:125209. doi:10.1016/j.envpol.2024.125209
174. Luo T, Pan Y, Liu Y, et al. LANA regulates miR-155/GATA3 signaling axis by enhancing c-Jun/c-Fos interaction to promote the proliferation and migration of KSHV-infected cells. *J Med Virol*. 2023;95(1):e28255. doi:10.1002/jmv.28255
175. Rastegar-Moghaddam SH, Ebrahimzadeh-Bideskan A, Shahba S, Malvandi AM, Mohammadipour A. Roles of the miR-155 in Neuroinflammation and Neurological Disorders: A Potent Biological and Therapeutic Target. *Cell Mol Neurobiol*. 2023;43(2):455-467. doi:10.1007/s10571-022-01200-z
176. Figlia G, Gerber D, Suter U. Myelination and mTOR. *Glia*. 2018;66(4):693-707. doi:10.1002/glia.23273
177. Minard AY, Tan SX, Yang P, et al. mTORC1 Is a Major Regulatory Node in the FGF21 Signaling Network in Adipocytes. *Cell Rep*. 2016;17(1):29-36. doi:10.1016/j.celrep.2016.08.086
178. Witt O, Sand K, Pekrun A. Butyrate-induced erythroid differentiation of human K562 leukemia cells involves inhibition of ERK and activation of p38 MAP kinase pathways. *Blood*. 2000;95(7):2391-2396. doi:10.1182/blood.V95.7.2391
179. Desideri E, Vegliante R, Cardaci S, Nepravishta R, Paci M, Ciriolo MR. MAPK14/p38α-dependent modulation of glucose metabolism affects ROS levels and autophagy during starvation. *Autophagy*. 2014;10(9):1652-1665. doi:10.4161/auto.29456
180. Asih PR, Prikas E, Stefanoska K, Tan ARP, Ahel HI, Ittner A. Functions of p38 MAP Kinases in the Central Nervous System. *Front Mol Neurosci*. 2020;13:570586. doi:10.3389/fnmol.2020.570586
181. Canagarajah BJ, Khokhlatchev A, Cobb MH, Goldsmith EJ. Activation Mechanism of the MAP Kinase ERK2 by Dual Phosphorylation. *Cell*. 1997;90(5):859-869. doi:10.1016/S0092-8674(00)80351-7
182. Esteras N, Blacker TS, Zherebtsov EA, et al. Nrf2 regulates glucose uptake and metabolism in neurons and astrocytes. *Redox Biol*. 2023;62:102672. doi:10.1016/j.redox.2023.102672
183. Troy CM, Shelanski ML. Down-regulation of copper/zinc superoxide dismutase causes apoptotic death in PC12 neuronal cells. *Proc Natl Acad Sci*. 1994;91(14):6384-6387. doi:10.1073/pnas.91.14.6384
184. Philips T, Bento-Abreu A, Nonneman A, et al. Oligodendrocyte dysfunction in the pathogenesis of amyotrophic lateral sclerosis. *Brain*. 2013;136(2):471-482. doi:10.1093/brain/aws339
185. See WZC, Naidu R, Tang KS. Cellular and Molecular Events Leading to Paraquat-Induced Apoptosis: Mechanistic Insights into Parkinson's Disease Pathophysiology. *Mol Neurobiol*. 2022;59(6):3353-3369. doi:10.1007/s12035-022-02799-2
